# Rnf20 shapes the endothelial control of heart morphogenesis and function

**DOI:** 10.1101/2022.09.16.508288

**Authors:** Linda Kessler, Rui Gao, Nalan Tetik-Elsherbiny, Olga Lityagina, Azhar Zhailauova, Yonggang Ren, Felix A. Trogisch, Julio Cordero, Yanliang Dou, Yinuo Wang, Evgeny Chichelnitskiy, Joscha Alexander Kraske, Patricia Laura Schäfer, Chi-Chung Wu, Guillermo Barreto, Michael Potente, Thomas Wieland, Roxana Ola, Joerg Heineke, Gergana Dobreva

**Affiliations:** Department of Cardiovascular Genomics and Epigenomics, European Center for Angioscience (ECAS), Medical Faculty Mannheim, Heidelberg University, Mannheim, Germany; Max Planck Institute for Heart and Lung Research, Bad Nauheim, Germany; Department of Cardiovascular Physiology, ECAS, Medical Faculty Mannheim, Heidelberg University, Mannheim, Germany; Department of Medical Oncology, University Hospital Heidelberg (UKHD) and National Center for Tumor Diseases (NCT), Heidelberg, Germany; Department of Molecular and Radiooncology, German Cancer Research Center (DKFZ), Heidelberg, Germany; Ploidy and Organ Physiology Laboratory, ECAS, Heidelberg University, Mannheim, Germany; Université de Lorraine, CNRS, Laboratoire IMoPA, UMR 7365; Nancy, France.; Berlin Institute of Health at Charité—Universitätsmedizin Berlin, Berlin, Germany; Max Delbrück Center for Molecular Medicine in the Helmholtz Association, Berlin, Germany; Experimental Pharmacology, European Center for Angioscience, Medical Faculty Mannheim, Heidelberg University, Mannheim, Germany; Cardiovascular Pharmacology, European Center for Angioscience, Medical Faculty Mannheim, Heidelberg University, Mannheim, Germany.; German Centre for Cardiovascular Research (DZHK)

**Author notes:** Correspondence: Prof. Dr. Gergana Dobreva, Medical Faculty Mannheim/ University of Heidelberg, Ludolf-Krehl-Str. 7-11, 68167 Mannheim, Germany.

**Keywords:** endothelial-cardiomyocyte crosstalk, heart development, congenital heart disease, endothelial-to-mesenchymal transition (EndMT), Rnf20, H2B ubiquitination (H2Bub), cardiac progenitor cells, cardiomyocyte binucleation, premature cardiomyocyte maturation

## Abstract

During embryogenesis, distinct cardiac cell types form, which shape the structural and functional properties of the heart. How their activity is coordinated is largely unknown. Here we show that Rnf20 is a multifaceted regulator of cardiac morphogenesis and function. On the one hand, Rnf20 controls extracellular matrix dynamics and endothelial-cardiomyocyte crosstalk essential for second heart field development. On the other hand, it safeguards endothelial cell identity and function by maintaining physiological angiocrine signaling and preventing endothelial-to-mesenchymal transition. Endothelial-specific deletion of Rnf20 led to ventricular septal defects, myocardial thinning and cardiac dysfunction as a result of aberrant signaling and excessive extracellular matrix deposition that induced precocious cardiomyocyte binucleation and irregular contractility. Furthermore, we uncovered upstream factors (e.g. Sox9) and multiple angiocrine and extracellular matrix molecules that alter cardiomyocyte functionality upon endothelial Rnf20 loss. In summary, our work identifies a novel, endothelial-specific role of Rnf20 in regulating cardiac morphogenesis and function.

## Introduction

The heart is a complex organ composed of distinct cell types, which shape its structural, mechanical and electrical properties (Meilhac and Buckingham, 2018). In particular, the crosstalk between endothelial cells and cardiomyocytes plays a crucial role both for normal cardiac morphogenesis and in cardiac disease. However little is known about the molecular players safeguarding physiological signaling between these cell types. During embryogenesis, endocardial endothelial cells (ECs) and myocardial precursors arise bilaterally within the mesoderm and through migration and embryo folding coalesce at the ventral midline to form a primitive heart tube consisting of an inner endocardial layer and an outer myocardial layer (Feulner et al., 2022; Tian and Morrisey, 2012). The two layers are separated by a dense layer of extracellular matrix (ECM), termed cardiac jelly, but communicate through paracrine signals. After formation of the initial heart tube, the heart grows by the addition of second heart field (SHF) progenitor cells to its anterior and venous poles (Cai et al., 2003; Gao et al., 2019; Meilhac and Buckingham, 2018; Vincent and Buckingham, 2010; Witzel et al., 2012). These SHF precursors, marked by the expression of the transcription factor (TF) Isl1, are multipotent and give rise to the distinct lineages in the heart (Caputo et al., 2015; Moretti et al., 2006; Zhuang et al., 2013), including myocardial and endocardial cells. The endocardium has recently gained attention due to its remarkable cellular plasticity, as well as its important signaling function during heart development, disease, and regeneration (Zhang et al., 2018). On one hand, it forms a vascular endothelial layer contiguous with the rest of the vasculature, which provides nutrients and oxygen to the heart. On the other hand, it receives and secretes signals that instruct heart morphogenesis and function, e.g. trabeculation, cardiac valve and septum formation, and myocardial compaction, thereby functioning as a crucial signaling interface (Feulner et al., 2022; Kim et al., 2021; Tian and Morrisey, 2012). With the formation of the compact myocardium intramyocardial vessels develop, which also signal to the subjacent cardiomyocytes (CMs) to regulate cardiac growth, contractility and rhythmicity. The importance of ECs for heart development has been demonstrated by endothelial-specific gene inactivation of different components of the Notch, vascular endothelial growth factor (VEGF), bone morphogenic protein (BMP), transforming growth factor (TGF), fibroblast growth factor (FGF), Neuregulin-1 (Nrg1)/Erbb2/b4 and angiopoietin-1 signaling pathways (Feulner et al., 2022; Kim et al., 2021; Tian and Morrisey, 2012). However, how signal-dependent transcription is coordinated during heart development is largely unknown.

At the molecular level, these dynamic cell-cell interactions induce signaling cascades, which converge in the nucleus to induce transcriptional and epigenetic programming that directs cell fate and function (Elsherbiny and Dobreva, 2021), while at the same time allowing a certain level of cell plasticity that is critical for proper cardiac morphogenesis and tissue healing following environmental stress. The classical model of inducible gene expression is based on signal-dependent RNA polymerase II (Pol II) recruitment and transcription initiation by inducible binding of transcription factors and epigenetic modifiers to promoters and enhancers. However, more recent studies have shown that inducible transcription is often regulated after transcription initiation, by pausing and release of promoter-proximal Pol II (Hargreaves et al., 2009; Levine, 2011; Liu et al., 2015). Many factors have been identified to regulate Pol II pausing and elongation, including Rnf20, the major E3 ubiquitin ligase responsible for monoubiquitination of histone H2B at lysine 120 (H2BK120ub, H2Bub11) (Kim et al., 2009; Kim et al., 2005; Pavri et al., 2006). Mechanistically, Rnf20 plays a key role in transcriptional control, by altering nucleosome stability and acting as a signaling hub for downstream processes associated with transcript pausing and elongation by Pol II, such as co-transcriptional mRNA processing (Chen et al., 2015; Fuchs and Oren, 2014; Pavri et al., 2006; Shema et al., 2011). The important function of Rnf20 for proper heart development was revealed by exome sequencing data, which identified a *de novo* mutation in *RNF20* to be associated with congenital heart defects in human patients (Zaidi et al., 2013). Further work in Xenopus found that *Rnf20* depletion results in defects in left-right (LR) asymmetry likely due to impaired cilia motility (Robson et al., 2019), while a recent study revealed a critical role of Rnf20 in CMs for CM maturation in the early postnatal window (VanDusen et al., 2021). Despite these interesting findings, the role of Rnf20 in the different cardiac cell types for ensuring proper cardiogenesis is largely unexplored.

In this study, we have addressed the role of Rnf20 in heart development. We find that Rnf20 is highly expressed in SHF progenitors and orchestrate SHF morphogenesis. Rnf20 ablation in Isl1 positive (Isl1+) cardiovascular progenitors and their progeny results in severe cardiac and vascular abnormalities, including shorter outflow tract (OFT), hypoplastic ventricular wall, abnormal trabeculation and disorganized endocardium. Functional and molecular analyses revealed that Rnf20 plays a central role in cardiac ECs for proper cardiac morphogenesis and function by inhibiting endothelial-to-mesenchymal transition (EndMT) and aberrant angiocrine signaling that induce CM cell cycle withdrawal and arrhythmic beating behavior.

## Results

### *Rnf20* is crucial for SHF development

The transcription factor Isl1 orchestrates a complex gene regulatory network driving SHF development and *Isl1* ablation results in loss of SHF derivatives, including the right ventricle (RV), OFT and large portions of the atria (Cai et al., 2003; Caputo et al., 2015; Gao et al., 2019). To further unravel the molecular determinants of Isl1-mediated SHF development, we performed a protein array screen for protein interaction partners of Isl1 and identified Rnf20, an E3 H2Bub1 ligase, which tightly associates with Pol II. Co-immunoprecipitation experiments in cardiac progenitors confirmed the interaction, while GST pulldown assays further validated that the binding of Isl1 and Rnf20 is direct, suggesting that Isl1 might regulate transcription through its direct interaction with Rnf20 (Figure 1A). Analysis of single cell RNA-Sequencing (RNA-Seq) datasets of E8.25 embryos (Xiong et al., 2019) showed higher expression of Rnf20 in SHF progenitors, further supporting a role of Rnf20 in Isl1-driven SHF development (Figure 1B). To understand the function of Rnf20 in the developing heart, we first conditionally ablated Rnf20 in mice using *Isl1-Cre*, which deletes in Isl1+ cardiovascular progenitors and their downstream progeny, i.e. the myocardium and the endocardium. *Rnf20*-deficient embryos died by E14.5 with severe cardiac and vascular abnormalities, including shorter OFT, hypoplastic ventricular wall, abnormal trabeculation and disorganized endocardium (Figure 1C-G). Co-immunostaining for the proliferation marker EdU together with isolectin B4 (IB4), an endothelial marker, and myosin heavy chain (MF20), a CM marker, revealed significantly decreased CM proliferation in the right and the left ventricle (LV) of E12.5 *Rnf20*-deficient hearts (Figure 1H, 1I). Immunostaining for H2Bub1, a histone modification catalyzed by Rnf20, revealed complete loss of H2Bub1 in the RV, whereas CMs in the LV were positive for H2Bub1 upon Rnf20-depletion using the Isl1-Cre deleter (Figure 1J, 1K). Interestingly, the endocardial cells in the RV, as well as most of those in the LV, were negative for H2Bub1 (Figure 1K), suggesting that Rnf20 might play a role in endocardial cells, which is important for CM proliferation.

**Figure 1.**
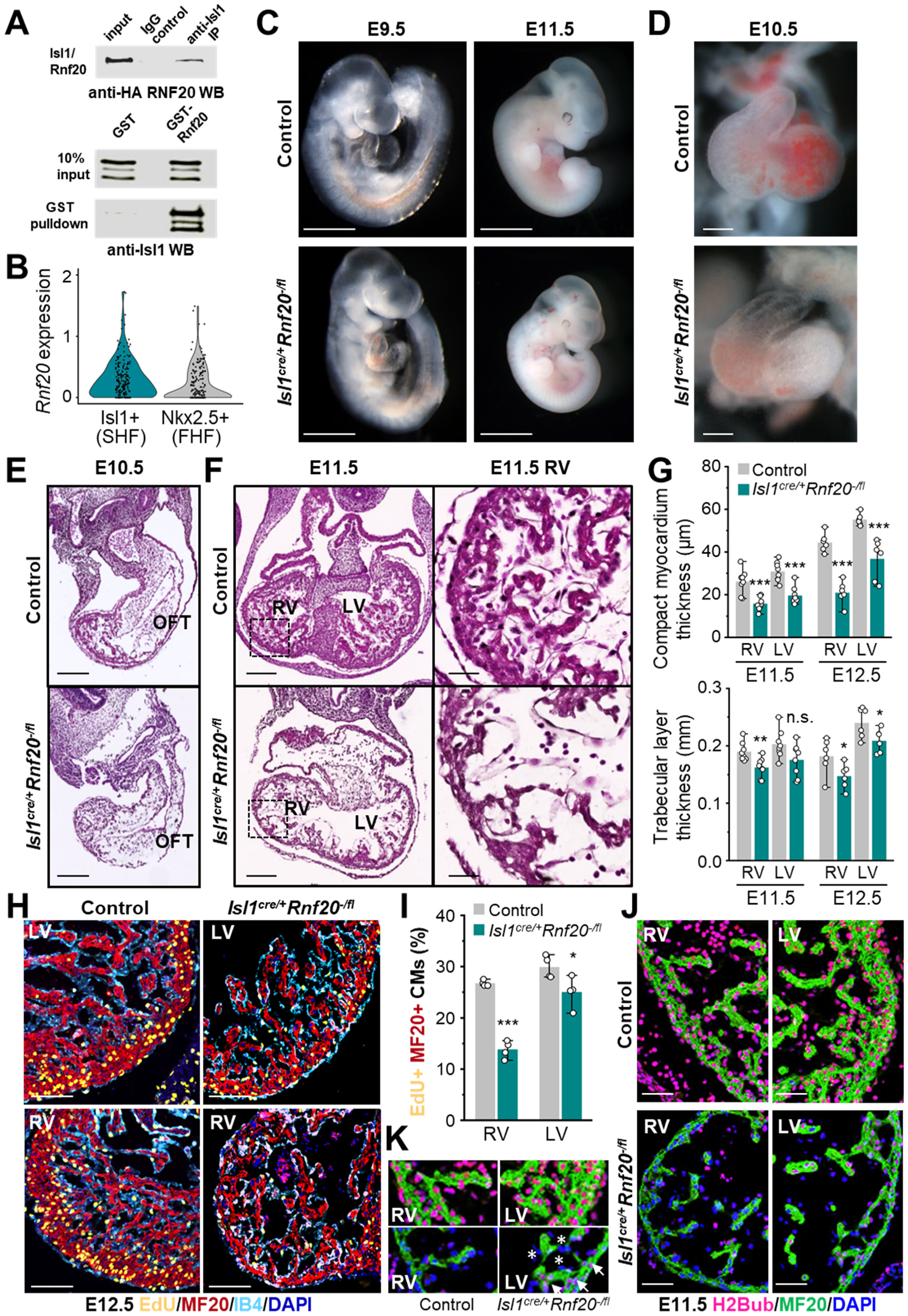
Ablation of Rnf20 in Isl1+ cardiovascular progenitors results in defects in SHF development. (A) Co-immunoprecipitation showing interaction between Isl1 and Rnf20 (top). GST pull down assays with recombinant GST protein or GST-Rnf20 fusion protein and recombinant Isl1 protein, showing that the interaction between those two proteins is direct (bottom). (B) Violin plot of *Rnf20* expression levels in Isl1+ and NKX2.5+ single cells of E8.25 embryos (Xiong et al., 2019). Each black dot represents a single cell. FHF, First heart field. SHF, Second heart field. (C) Gross appearance of control (*Isl1^+/+^Rnf20^+/fl^*) and *Isl1^cre/+^Rnf20^-/fl^* embryos at E9.5 and E11.5. Scale bars, 2 mm (left), 1 mm (right). (D) Frontal views of E10.5 control and *Isl1^cre/+^Rnf20^-/fl^* hearts, showing short outflow tract (OFT) and a smaller right ventricle (RV). Scale bars, 100 µm. (E) H&E stainings of transverse sections of control and *Isl1^cre/+^Rnf20^-/fl^* embryos at E10.5 showing shortening of the OFT. Scale bars, 200 µm. (F) Histological analysis of control and Rnf20 knockout hearts at E11.5 showing a smaller RV, hypoplastic ventricular wall and abnormal trabeculation. Magnified images of indicated RV regions (square) are shown in the right panel. Scale bars, 200 µm (left), 50 µm (right). (G) RV and left ventricle (LV) wall thickness (top) and trabecular layer thickness (bottom) in control (n=8 for E11.5, n=6 for E12.5) and *Isl1^cre/+^Rnf20^-/fl^* embryos (n=6 for E11.5, n=5 for E12.5) of E11.5 and E12.5 hearts. (H, I) Immunostaining with EdU, anti-MF20 (CMs), anti-IB4 (ECs) and DAPI (nucleus) (H) and quantification of EdU-labeled CMs (EdU+/MF20+ cells) in RV and LV of E12.5 control and *Isl1^cre/+^Rnf20^-/fl^* embryos (n=4) (I), showing a marked reduction in the number of proliferating CMs upon Rnf20 ablation. Scale bars, 100 µm. (J, K) Immunostaining for H2Bub1 catalyzed by Rnf20 and MF20 in RV and LV of E11.5 control and *Isl1^cre/+^*-Rnf20^-/fl^ embryos (J) and magnified areas (K). Arrows point to H2Bub1-positive CMs in the LV; asterisks indicate H2Bub1-negative endocardial cells. Scale bars, 50 µm. For this and all other figures, error bars represent mean ±SD. *p<0.05 **p<0.01 ***p<0.001.

### Rnf20 controls extracellular matrix dynamics and cell signaling essential for SHF development

To elucidate Rnf20’s downstream effectors governing cardiogenesis, we performed RNA-Seq of OFT and RV dissected from E9.5 and E10.5 control and *Isl1^cre/+^*Rnf20^-/fl^ embryos (Figure 2A-C, Figure S1A, Table S1). Gene Ontology (GO) pathway analysis of E9.5 hearts revealed an over-representation for genes linked to cell migration and adhesion, ECM organization, angiogenesis as well as transcriptional regulation, including TFs involved in EndMT such as Snai1, Snai2 and Msx2. In contrast, genes downregulated upon Rnf20 loss of function (LOF) were linked to cell cycle, Notch signalling and muscle contraction (Figure 2C). However, at E10.5, genes involved in blood vessel and heart development as well as cell-cell signalling were significantly downregulated upon Rnf20 LOF, whereas upregulated genes upon Rnf20 loss were mainly involved in sarcomere organization, regulation of muscle contraction and similarly cell-cell signalling (Figure 2D, Figure S1B), suggesting a dynamic function of Rnf20 during cardiogenesis. In line with the known function of Rnf20 in transcriptional elongation, differentially expressed genes upon Rnf20 loss have a low pausing index (Figure S1D-F). Since Rnf20 interacts with Isl1, which induces chromatin opening of key SHF genes (Gao et al., 2019), we next analysed chromatin accessibility at Rnf20 target genes. We found a major decrease of chromatin accessibility at Rnf20 targets in Isl1-/- mice (Figure S1C), supporting the notion that Isl1 and Rnf20 work together to instruct SHF morphogenesis. Furthermore, we observed a significant overlap between Rnf20 targets and genes deregulated in dissected OFT and RV of E10.5 *Isl1* haploinsufficient mice (Gao et al., 2019), particularly for ECM genes and a number of proteases of the Adamts family (A disintegrin and metallopeptidase with thrombospondin motif), i.e. Adamts2/3/10/20 and matrix metalloproteinases (Mmps) (Figure 2E).

**Figure 2.**
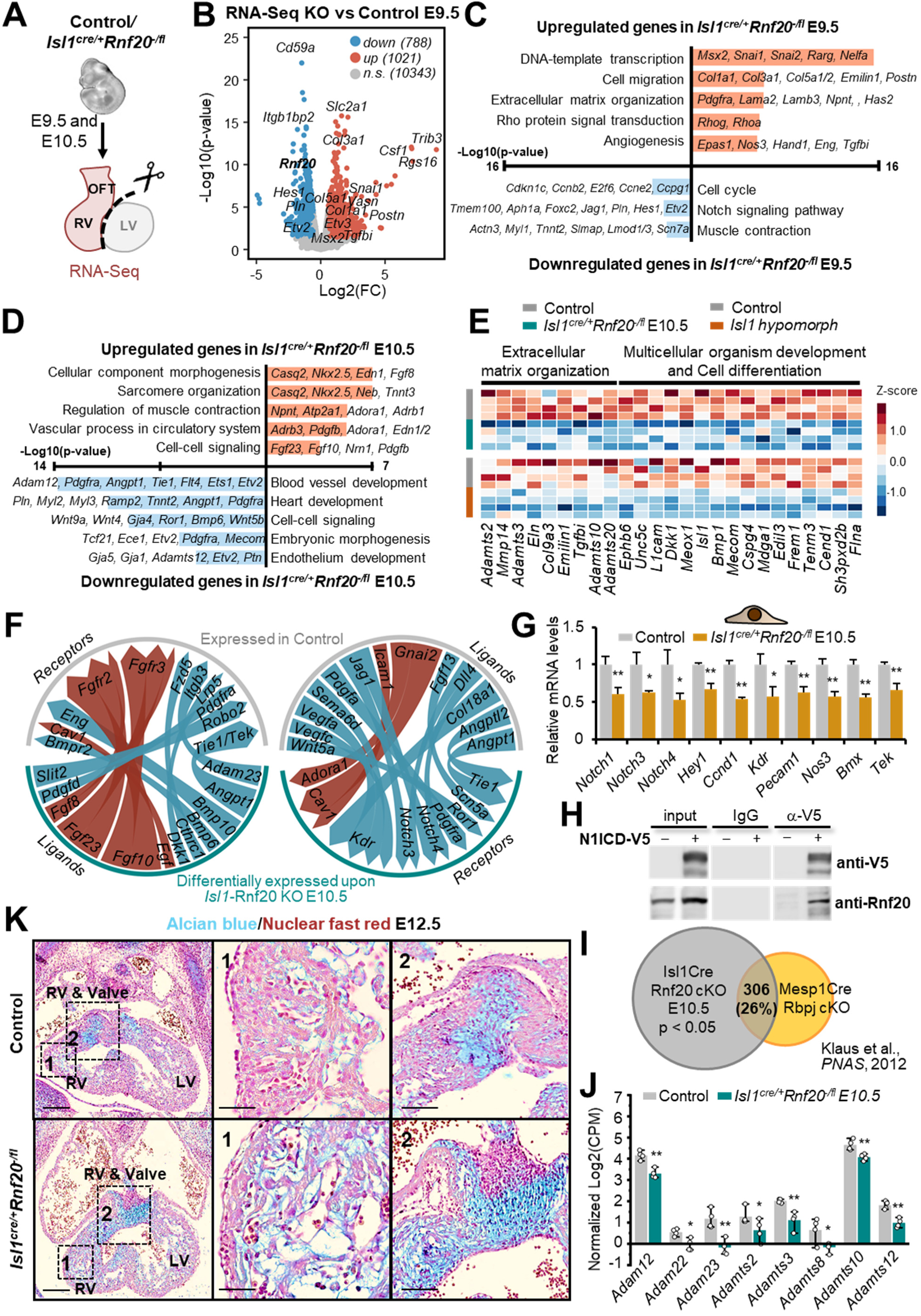
Rnf20 controls ECM dynamics and signaling in SHF development. (A) Schematic representation of the dissection procedure used for RNA-Seq experiments. RV and OFT of E9.5 and E10.5 control and *Isl1^cre/+^Rnf20^-/fl^* were dissected. (B) Volcano plot showing the distribution of differentially expressed genes in dissected E9.5 OFT and RV of *Isl1^cre/+^Rnf20^-/fl^* versus control OFT and RV (n=4; Log2(FC) ≤ -0.58, ≥0.58; p-value < 0.05). Representative up- (red) and downregulated (blue) genes are indicated. (C) Gene ontology pathway enrichment analysis and representative up- (red) and downregulated (blue) genes in E9.5 *Isl1^cre/+^Rnf20^-/fl^*compared to control hearts. (n=3; Log2(FC) ≤ -0.58, ≥0.58; p-value < 0.05). (D) Gene ontology pathway enrichment analysis and representative up- (red) and downregulated (blue) genes in E10.5 *Isl1^cre/+^Rnf20^-/fl^* compared to control hearts. (E) Heatmap representation of genes involved in extracellular matrix organization and cell differentiation in dissected OFT and RV of Rnf20-deficient and Isl1 haploinsufficient hearts (Gao et al., 2019). (F) Circos plot of the significantly changed ligand-receptor interactions at E10.5 upon Rnf20 LOF in Isl1+ cardiovascular precursors. Red arrows represent upregulated ligands, blue arrows represent downregulated ligands and arrow thickness indicates the probability for interaction, i.e. interaction score (Skelly et al., 2018). (G) Relative mRNA expression levels of genes involved in endothelial-myocardial interactions in ECs isolated from control and *Rnf20* KO E10.5 RV and OFT (n= 3). (H) Co-immunoprecipitation with N1ICD and Rnf20, showing that Rnf20 interacts with N1ICD. (I) Overlap between genes differentially expressed in Rnf20 and Rbpj knockout (Klaus et al., 2012) compared to control hearts. (J) Relative expression of genes associated with cardiac jelly protein degradation in control and *Isl1^cre/+^Rnf20^-/fl^* RV and OFT. (K) Images of Alcian blue and nuclear fast red staining of transverse sections of control and *Isl1^cre/+^Rnf20^-/fl^* hearts at E12.5. Magnified images of indicated regions are shown in the right panels. Scale bars, 200 µm (whole heart), 50 µm (magnified regions).

Since genes involved in cell-cell signalling were deregulated upon Rnf20 depletion, we next performed ligand-receptor analysis (Figure 2F, Table S2). Among the downregulated ligands we found *Angpt1* and *Bmp10*, which play key roles in chamber morphogenesis, trabeculation and myocardial compaction, while among the downregulated receptors we uncovered *Tie1*, *Pdgfra*, *Kdr*, *Notch3* and *Notch4* (Figure 2F). Importantly, genes involved in endocardial-myocardial interactions, e.g. Vegfr2 (Kdr), Notch1/2/3 and the Notch target genes Hey1, Hes1, Erbb2 and Ccnd1 were significantly downregulated in endocardial cells isolated from Rnf20 deficient E10.5 hearts (Figure 2G). Rnf20 has been shown to be stimulate Notch reporter gene expression and Rnf20 loss of function resulted in Notch signaling defects in Drosophila (Bray et al., 2005). Therefore, we next studied whether Notch1 and Rnf20 interact. Indeed, co-immunoprecipitation analysis revealed that the active Notch1 intracellular domain (N1ICD) interacts with Rnf20 (Figure 2H). Further, by intersecting transcriptomics datasets of hearts mutant for the principal transcriptional mediator of Notch signalling, Rbpj (Klaus et al., 2012; MacGrogan et al., 2018), and for Rnf20 we observed a significant overlap (Figure 2I), suggesting that Rnf20 works together with Notch1 to regulate cardiac morphogenesis. Notch1 controls cardiac jelly dynamics through regulating *Adamts1* and *Mmp2* levels (Del Monte-Nieto et al., 2018). *Mmp2, Mmp14* and a number of proteases of Adam (A disintegrin and metalloproteinases) and Adamts (A disintegrin and metallopeptidase with thrombospondin motif) families were also significantly downregulated upon Rnf20 LOF (Figure 2J). Consistent with the reduced expression of *Mmps* and *Adam*- and *Adamts*- family members, alcian blue staining revealed excessive accumulation of cardiac jelly upon *Rnf20* LOF, particularly around the ventricular trabeculae and the endocardial cushions (Figure 2K).

### Rnf20 plays a key role in cardiac ECs for heart development and function

Rnf20 has been shown to regulate CM maturation after birth (VanDusen et al., 2021), however its function in cardiac ECs is largely unknown. Interestingly, H2Bub1 was depleted in the RV and the majority of the LV endocardium, but was absent only in right ventricular CMs upon *Rnf20* ablation in SHF progenitors using the Isl1-Cre line, while CM proliferation was significantly decreased in both ventricles. Moreover, we identified many critical pathways involved in EC-CM communications to be significantly deregulated upon Rnf20 LOF. These observations suggest that Rnf20 might play an important role in ECs for cardiac morphogenesis, which prompted us to further examine Rnf20 function in cardiac ECs. To this end, we inactivated Rnf20 using a *Tie2-Cre* deleter, which recombines in ECs and some hematopoietic cells (Koni et al., 2001). *Tie2-Cre*-mediated *Rnf20* deletion resulted in early embryonic lethality around E10.5 (Figure 3A). *Tie2^Cre^Rnf20^fl/fl^* mutant embryos displayed shortened OFT and small RV – structures generated by Isl1+ progenitor cells (Figure 3B), supporting the notion that endothelial Rnf20 is important for SHF development.

**Figure 3.**
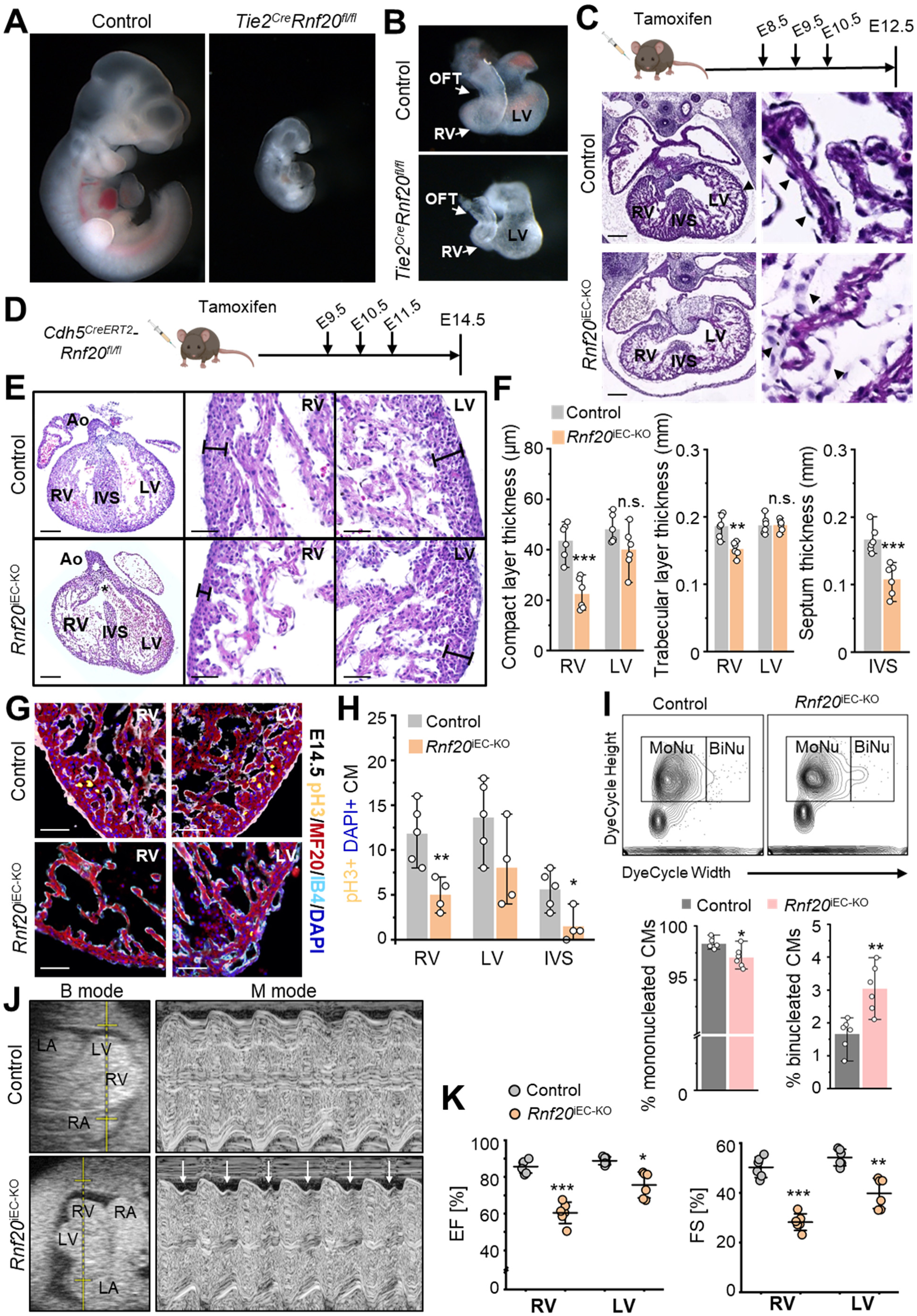
Endothelial Rnf20 is essential for heart development and function. (A, B) Gross appearance of control (*Tie2^Cre^Rnf20^+/fl^*) and *Tie2^Cre^Rnf20^fl/fl^* embryos at E10.5 (A) and of dissected hearts (B). (C) Schematic representation of the experimental setup and histological analysis of control and *Rnf20*^iEC-KO^ hearts at E12.5 showing a thinner compact layer, less developed trabeculae, disorganized interventricular septum (left) and disorganized endocardial cells (arrows in the right panel). Scale bars, 200 µm. (D) Schematic representation of the experimental setup. For this experiment and all other studies, control (*Cdh5^CreERT2^*neg-*Rnf20^fl/fl^*) and *Rnf20*^iEC-KO^ (*Cdh5^CreERT2^*pos-*Rnf20^fl/fl^*) embryos were exposed to tamoxifen from E9.5. (E) H&E staining of representative paraffin sections of E14.5 hearts of control and *Rnf20*^iEC-KO^ embryos (left panels), and higher magnification of RVs and LVs (boxed area) showing thinner compact myocardium and trabecular layer (middle and right panels). Abbreviations: Ao, Aorta; LV, left ventricle; RV, right ventricle; IVS, interventricular septum. Star indicates ventricular septal defect (VSD). Scale bars, 200 µm (whole heart), 50 µm (magnified regions). (F) Quantification of the thickness of the compact and trabecular layer as well as the ventricular septum thickness in control and *Rnf20*^iEC-KO^ hearts (n=6). (G, H) Immunostaining of E14.5 control (n=5) and *Rnf20*^iEC-KO^ (n=4) heart sections with Ser10 phospho-H3 (pH3+) and MF20 antibody (G) and quantification of the percentage of mitotic pH3+ CMs (H). Scale bars, 50 µm. (I) Representative FACS analysis of staining with Vybrant DyeCycle DNA dye for mono- and binucleation in *Rnf20*^iEC-KO^ hearts (top panel). The percentage of mononucleated and binucleated CMs (n=6, bottom panel). (J) Representative examples of B- and M-mode echocardiograms of control and *Rnf20*^iEC-KO^ embryos at E14.5, showing irregular contractility of the RV. (K) Quantification of the ejection fraction (EF) and fractional shortening (FS) for RV and LV assessed by echocardiography (n=6).

To be able to study the role of endothelial Rnf20 at different embryonic stages we next used the tamoxifen-inducible *Cdh5-CreERT2* and *Pdgfb-CreERT2* endothelial-targeting lines (Claxton et al., 2008; Wang et al., 2010) to induce Rnf20 ablation, which resulted in similar phenotypes (Figure 3C-K, Figure S2, data not shown). Endothelial deletion of Rnf20 in ECs following administration of tamoxifen from E8.5 resulted in thinner compact layer, fewer or less-developed trabeculae and a disorganized interventricular septum (IVS) at E12.5 (Figure 3C, Figure S2A) and lethality by E14.5. Interestingly, in contrast to wild-type embryos, in which endocardial cells typically show elongated nuclei and thin cell bodies lining the myocardium, Rnf20-depleted endocardial cells appeared disorganized and did not line the trabeculae (Figure 3C). Tamoxifen administration from E9.5 onwards resulted in similar cardiovascular abnormalities, i.e. reduced trabecular myocardium, thinner compact myocardium and reduced IVS wall thickness together with ventricular septal defects (VSD) (Figure 3D-F, Figure S2B, S2C). We next studied whether the observed alterations are the result of altered CM proliferation. Indeed, immunostaining for the mitotic marker Ser10 phosphorylated-histone H3 (pH3) in combination with the CM marker MF20 revealed a significant decrease in CM proliferation (Figure 3G, 3H). Furthermore, stainings with Vybrant DyeCycle DNA dye followed by FACS analysis, which can be used to distinguish between mononucleated and binucleated CMs (Windmueller et al., 2020), found significant increase in binucleated CMs in *Rnf20*^iEC-KO^ hearts (Figure 3I). Further, echocardiography measurements revealed irregular contractility of the RV as well as reduced ejection fraction, fractional shortening and cardiac output in *Rnf20*^iEC-KO^ embryos, indicating cardiac dysfunction (Figure 3J, 3K, Figure S2D).

### Direct co-differentiation of endocardial cells and CMs from pluripotent stem cells reveals perfusion independent effects of endothelial Rnf20 deletion on CMs

To study the role of Rnf20 in ECs in controlling CM behavior without the potential confounding side effects of altered tissue perfusion in malformed hearts, we established protocols for directed differentiation of murine ESCs (mESCs) in Nkx2.5+/CD31+ endocardial cells (based on (Mikryukov et al., 2021)), as well as protocols for co-differentiation of Nkx2.5+/CD31+ endocardial cells and Nkx2.5+/CD31- CMs from common cardiovascular progenitors (Figure 4A-F). Both co-differentiation and co-culture of CMs with ESC-derived endocardial ECs resulted in increased CM contractility and differentiation, as well as the expression of genes involved in EC-CM crosstalk (Figure 4D-H), suggesting that co-differentiation is a suitable system to study the CM-EC crosstalk during cardiogenesis. In contrast, co-culture of CMs with hemogenic ECs (Nkx2.5-/CD31+) did not significantly alter CM contractility. We then established induced pluripotent stem cells (iPSCs) from control (*Cdh5*^CreERT2^-negative-Rnf20^fl/fl^) and *Rnf20*^iEC-KO^ (*Cdh5*^CreERT2^-positive-Rnf20^fl/fl^) fibroblasts, which allow us to ablate Rnf20 specifically in ECs at distinct steps during cardiovascular differentiation using 4-hydroxytamoxifen (4-OHT) (Figure 4I). Adding 4-OHT to the differentiation medium at day 5 (cardiovascular progenitors) to induce Rnf20 ablation resulted in decreased CM proliferation and increased number of binucleated CMs, while proliferation of ECs was also reduced (Figure 4J-L). Analysis of movies of spontaneously beating CMs using MYOCYTER (Grune et al., 2019) revealed arrhythmicity (Figure 4M), similar to the phenotype observed *in vivo*. These findings suggest that the observed *in vivo* phenotypes are a direct consequence of Rnf20 ablation.

**Figure 4.**
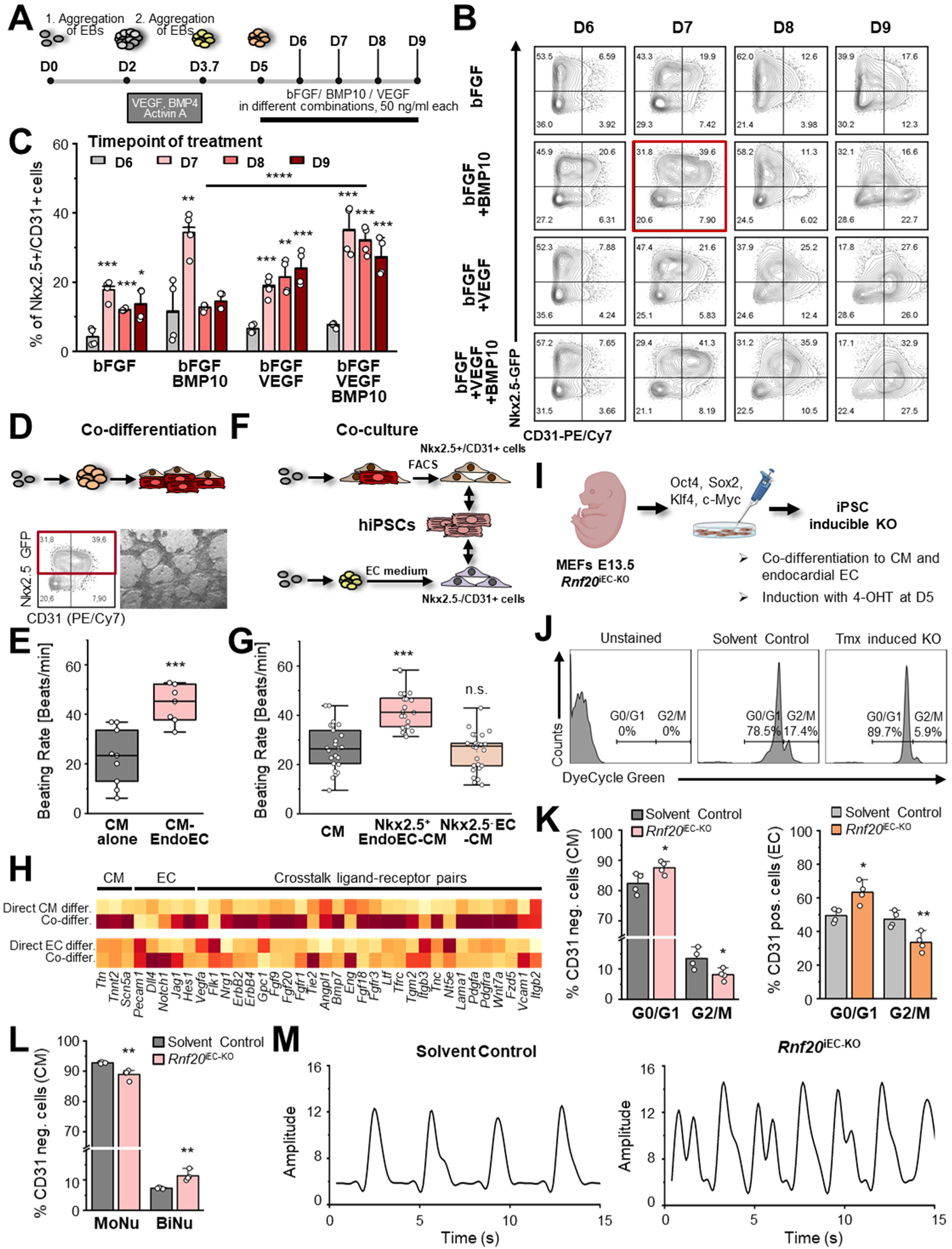
Endothelial Rnf20 instructs CM behavior in an *in vitro* co-differentiation system. (A) Schematic representation of the experimental design for the establishment of a co-differentiation system from murine ESC to endocardial cells and cardiomyocytes. (B, C) Representative FACS analysis of direct Nkx2.5-GFP fluorescence and staining for the EC marker CD31 (B) and percentage of Nkx2.5+/CD31+ endocardial cells (n=4) (C). (D) Characterization of simultaneous differentiation of CMs and endocardial cells from ESCs (co-differentiation, top). FACS analysis showing similar percentage of Nkx2.5+/CD31+ endocardial cells and Nkx2.5+/CD31-CMs co-differentiated from ESCs (bottom left) and images of co-differentiated cells (bottom right). (E) Beating rate of CMs differentiated by the direct differentiation protocol of ESCs into CMs or co-differentiated with endocardial cells at day 9. Beating rate was extracted from video sequences using MYOCYTER. (F) Co-culture of Nkx2.5+/CD31+ endocardial cells and Nkx2.5-/CD31+ hemogenic ECs with human iPSC-derived CMs. Nkx2.5+/CD31+ endocardial cells were sorted from day 7 and differentiated using the protocol represented in panel A, while Nkx2.5-/CD31+ hemogenic ECs were differentiated by the addition of EC differentiation medium at mesoderm stage (protocol adapted for mouse ESCs from (Palpant et al., 2017). (G) Beating rate of hiPSC-CMs co-cultured either with Nkx2.5+/CD31+ endocardial cells or Nkx2.5-/CD31+ hemogenic ECs for 2 days. Beating rate was extracted from video sequences using MYOCYTER. (H) Relative expression of CM, EC marker genes as well as genes involved in EC-CM crosstalk in CMs differentiated by the direct differentiation protocol of ESCs into CMs or co-differentiated with endocardial cells (top) or ECs differentiated by the direct differentiation protocol of ESCs into ECs or co-differentiated with CMs (bottom). (I) Schematic representation of the methodology used for the generation of control and *Rnf20*^iEC-KO^ mouse iPSC lines. (J) Representative FACS analysis of 38 days old CMs (CD31 negative cells in the co-differentiation) stained with Vybrant DyeCycle DNA dye, showing decrease in CMs in G2/M phase upon endothelial Rnf20 ablation. For this experiment as well as the data shown in (K-M), control and *Rnf20*^iEC-KO^ iPSCs were co-differentiated in CM and endothelial specific Rnf20 ablation was induced at day 5 by 4-hydroxytamoxifen (4-OHT). (K) Quantification of the percentage of CMs (left) or endocardial cells (right) in G0/G1 and G2/M phase upon EC-specific tamoxifen-mediated Rnf20 ablation (n=4). Cells were differentiated using the co-differentiation protocol and stained with Vybrant DyeCycle DNA dye, and CD31, followed by FACS analysis. (L) Quantification of the percentage of mononucleated and binucleated CMs (day 38) of cells stained with Vybrant DyeCycle DNA dye and subjected to FACS analysis (n=4). (M) Plot of contraction amplitude and speed in spontaneously beating CMs (day 18) from video sequences using MYOCYTER.

### Rnf20 inhibits endothelial-to-mesenchymal transition (EndMT)

To further understand the mechanisms underlying the impact of endothelial Rnf20 on CMs, we performed single-cell RNA-Seq of E14.5 control and *Rnf20*^iEC-KO^ hearts as well as bulk RNA-Seq of isolated cardiac ECs and CMs (Figure 5A, Figure S3A-C). Similar to previous studies, we detected several distinct cell types with transcriptional signatures of different CM subtypes, endocardial cells, cardiac cushions, valves, vascular ECs, hemogenic endocardium, smooth muscle cells and others (Xiao et al., 2018) (Figure 5B). Consistent with our histological studies, single-cell RNA-Seq analysis revealed major changes in ventricular CM number, cell cycle activity determined by the expression of Mki67 (Marker of Proliferation Ki-67), and maturation as defined by the expression of mitochondrial marker genes such as mt-Co2, mt-Nd2, mt-Nd1 and mt-Atp6 (Figure 5B, 5C). We next re-clustered the EC populations to investigate alterations in Rnf20-deficient ECs in greater detail. This approach identified seven clusters, including three distinct endocardial cell populations, a proliferative endocardial population, one coronary EC population, as well as endocardial valve and AV cushion cells (Figure 5D). The proliferative endocardial cluster, marked by the Mki67 (Marker of proliferation Ki-67), was strongly reduced in Rnf20-deficient hearts (Figure 5D, 5E), consistent with the reduced endocardial cell proliferation in our *in vitro* co-differentiation system (Figure 4J, 4K). Interestingly, we observed many genes involved in ECM organization to be upregulated particularly in endocardial clusters, EndoC2+3 and endocardial valve cells (Figure 5E, 5F). Similarly, bulk RNA-Seq analysis revealed that genes upregulated in Rnf20-deficient ECs show over-representation of GO terms linked to connective tissue development and cell-cell signaling by Wnts and Bmps, while downregulated genes were related to chromatin modification, transcriptional activation, cell proliferation and communication (Figure 5G, Figure S3D). Gene set enrichment analysis further supported an important role of Rnf20 in the regulation of genes involved in cell-cell interactions (Figure S3E). Interestingly many typical markers of EndMT, e.g. fibronectin (*Fn1*), vimentin (*Vim*), *Icam1*, *Mmp2*, a large set of collagens as well as TFs driving EndMT in the heart such as *Hand2* and *Foxc1*, were highly upregulated upon *Rnf20* loss suggesting that Rnf20 safeguards EC identity by preventing EndMT (Figure 5G, Figure S3D, S3F). Immunostaining confirmed the high expression of Vim in *Rnf20*^iEC-KO^ endocardium (Figure 5H). Similarly, depletion of Rnf20 in HUVECs resulted in upregulation of genes involved in ECM organization and downregulation of genes regulating cell proliferation (Figure S3G). Further, Rnf20-deficient HUVECs showed increased migratory behavior (Figure S3H-J), supporting a function of Rnf20 in inhibiting EndMT.

**Figure 5.**
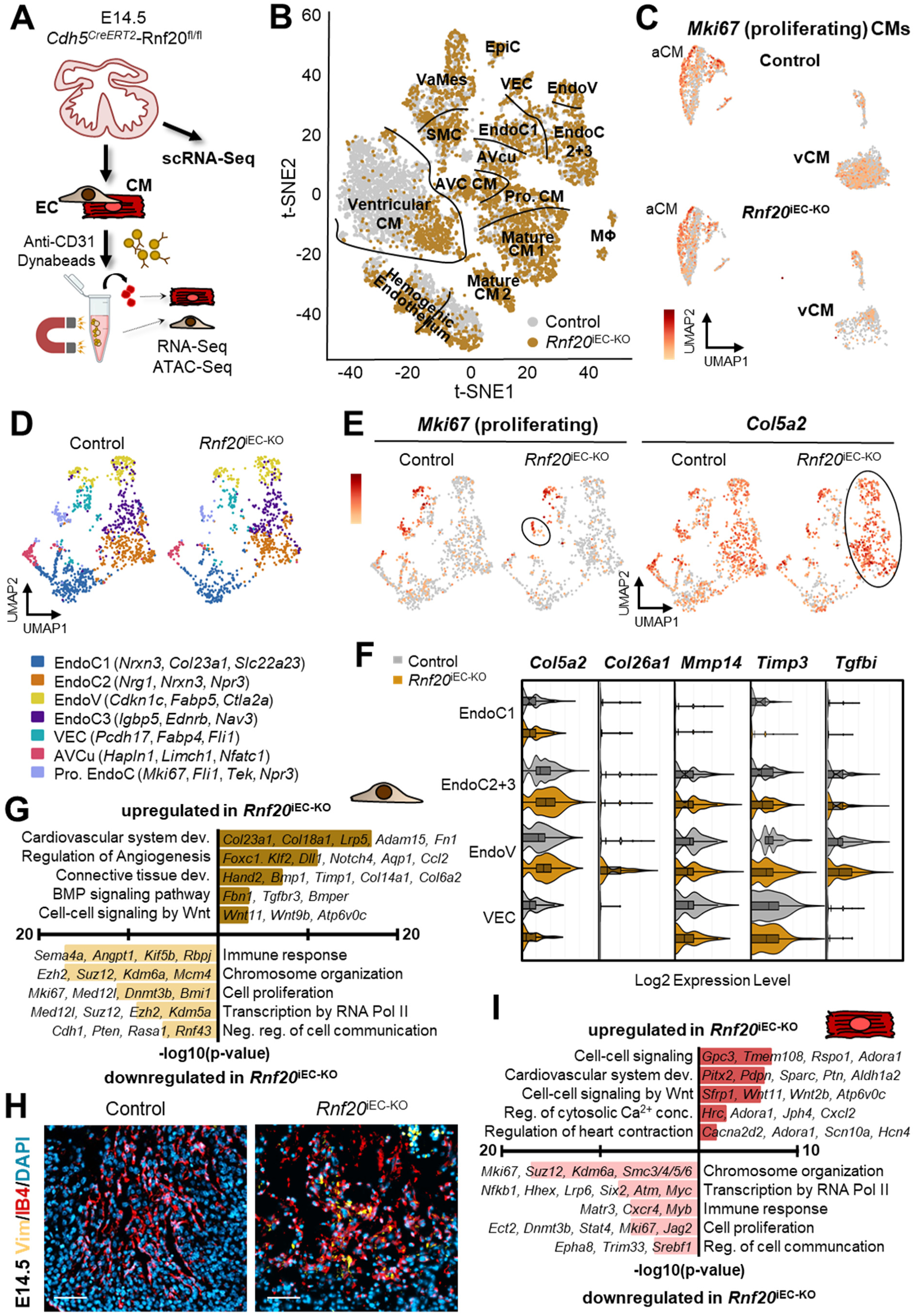
Endothelial Rnf20 inhibits EndMT, CM maturation and cell cycle withdrawal. (A) Schematic representation of the experimental setup for the analysis presented in Figure 5 and Figure 6. E14.5 control and *Rnf20*^iEC-KO^ hearts were used for scRNA-Seq and purified ECs and CMs for RNA-Seq and ATAC-Seq. (B) 10x genomics single-cell RNA-Seq of E14.5 control and *Rnf20*^iEC-KO^. AVC CM, atrioventricular cardiomyocytes (CM); AVcu, atrioventricular cushion cells; Pro. CM, proliferating CM; EndoC, endocardial cells; EndoV, endocardial valve cells; VaMes, valve mesenchyme; SMC, smooth muscle cells; VEC, vascular ECs; EpiC, epicardial cells; MΦ, macrophages. (C) UMAP analysis of CM populations from panel B. Feature plots visualizing *Mki67*-expressing proliferative CM. aCM, atrial CM; vCM, ventricular CM. (D) UMAP analysis of CD31+ endothelial and endocardial populations from panel B identifying 7 subclusters. (E) Feature plots of the expression of *Mki67* and *Col5a2*, showing a decrease in proliferative endocardial cells (circled population) and an increase in *Col5a2* expression in subset of endocardial ECs. (F) Violin plots visualizing expression levels of collagens and extracellular matrix proteins in different endothelial and endocardial populations. (G) Gene ontology pathway enrichment analysis and representative genes in upregulated and downregulated genes in *Rnf20*^iEC-KO^ ECs compared to ECs from control hearts. n=3; Log2(FC) ≤ -0.58, ≥0.58; p-value < 0.05. (H) Immunostaining of E14.5 control and *Rnf20*^iEC-KO^ heart sections with the EndMT marker vimentin (Vim), IB4 for ECs and DAPI for nuclei. Scale bars, 50 µm. (I) Gene ontology pathway enrichment analysis and representative genes in upregulated and downregulated genes in *Rnf20*^iEC-KO^ CMs compared to CMs from control hearts. n=3; Log2(FC) ≤ -0.58, ≥0.58; p-value < 0.05.

### Rnf20 controls cell-cell signaling-dependent transcription

Interestingly, we detected significant transcriptional changes in CMs upon Rnf20 deletion in ECs (Figure 5I, Figure S3D). Upregulated genes were enriched in GO terms linked to cell-cell signaling, regulation of Ca^2+^ concentration and heart contraction (Hcn4, Scn10a, Cacna2d2), consistent with the aberrant contractility observed in *Rnf20*^iEC-KO^ embryos. Similar to ECs, genes downregulated in CMs were linked to chromatin modification, transcriptional activation and cell proliferation (Figure 5I, Figure S3D). Among the genes downregulated upon Rnf20 depletion was the RhoGEF gene Ect2 (Figure S3D), which regulates cytokinesis. Loss of Ect2 function has been shown to induce CM binucleation and cell-cycle withdrawal, and is sufficient to suppress the CM proliferation during cardiac regeneration (Liu et al., 2019; Windmueller et al., 2020). Interestingly, we detected significant changes in Rnf20 levels in CMs upon Rnf20 deletion in ECs. Similarly, Rnf20 levels were significantly upregulated in CMs upon co-culture with ECs (Figure S3K).

To identify upstream regulatory factors driving cardiac dysfunction in *Rnf20*^iEC-KO^ embryos, we performed assay for transposase-accessible chromatin (ATAC) sequencing (ATAC-Seq) (Buenrostro et al., 2013) of ECs and CMs isolated from control and *Rnf20*^iEC-KO^ hearts. Endothelial depletion of Rnf20 resulted in major changes in chromatin accessibility in both ECs and CMs (Figure 6A, 6B, Figure S4A, 4B). Similar to the transcriptomics data, genes that gained chromatin accessibility in ECs were highly enriched in GO terms linked to cell-cell signaling and vasculature development, while genes that gained open chromatin in CMs were linked to heart development, Ca^2+^ homeostasis and cell-cell signaling (Figure 6B). Genes that lost chromatin accessibility in ECs were similarly linked to cell-cell signaling and specifically to Wnt signaling as well as embryonic development. In CMs, genes linked to cell differentiation, cardiac septum development and Wnt signaling showed reduced open chromatin (Figure 6B). Through integrative analysis, we identified genes whose transcriptional level positively correlate with chromatin accessibility at their respective genomic loci, indicating binding or loss of binding of transcriptional activator (Figure 6C, 6D, 6F, 6G, Figure S4C, S4D). We also observed genes whose transcriptional level inversely correlate with their chromatin accessibility, indicative of transcriptional repressor binding. We next performed Transcription factor Occupancy prediction By Investigation of ATAC-Seq Signal (TOBIAS) footprinting analysis (Bentsen et al., 2020a) to pinpoint transcriptional activators and repressors of Rnf20-dependent transcription. We identified multiple TFs, including Stat5, Pparg, Pax3 and Nkx2.5 etc. as potential transcriptional activators that are highly bound in *Rnf20*^iEC-KO^ ECs compared to control ECs, and Hes6, Msx1 and Bcl6 as potential repressors of Rnf20-dependent transcription (Figure S4E). Analogous analysis of *Rnf20*^iEC-KO^ CMs pointed to Pparg, Hand1, Tp53 as transcriptional activators engaged in CMs upon *Rnf20* LOF in ECs, while Tbx20, Irf8/9 as potential transcriptional repressors (Figure S4F). TFs binding to regulatory regions are highly dynamic; hence, differential TF binding does not necessarily result in altered chromatin accessibility. Therefore, we next examined the differential binding of TFs at all ATAC peaks in control and *Rnf20*^iEC-KO^ associated only to deregulated genes (Figure 6E, 6H). We found Foxc1, Hif1a, Smad4, Tbx2 and Pparg to bind highly at Rnf20-targets in *Rnf20*^iEC-KO^ ECs, while key EC identity TFs (Etv2, Fli1, Erg) and TFs involved in Notch signaling showed decreased binding (Figure 6E). Particularly, TFs involved in EndMT (Sox9, Hif1a, Tbx2/3, Smads) were highly bound at genes upregulated upon *Rnf20* LOF (Figure 6I). In CMs, tumor suppressors restricting cell proliferation (Tp53 and Tp73) were highly enriched at Rnf20 target genes, while E2 factor (E2F) family members showed lower binding, consistent with the decreased CM proliferation in *Rnf20*^iEC-KO^ hearts (Figure 6H). SOX9 overexpression is sufficient to activate mesenchymal genes in HUVECs and induce EndMT (Fuglerud et al., 2022), thus we next compared gene expression profiles of ECs overexpressing Sox9 and lacking Rnf20. We found a high degree of overlap (31%), particularly in genes upregulated in both cases (Figure 6J). Genes significantly upregulated upon Sox9 overexpression and *Rnf20* LOF belonged to GO terms including cell adhesion, ECM organization as well as genes involved in regulation of blood pressure (Figure 6K). Given the high degree of overlap, we next tested whether Sox9 upregulation could induce CM binucleation and thus cell cycle withdrawal. We observed significantly increased number of binucleated CMs upon co-culture with ECs overexpressing Sox9, suggesting that Sox9 might regulate an EndMT transcriptional program resulting in premature CM maturation upon *Rnf20* LOF (Figure 6L, 6M).

**Figure 6.**
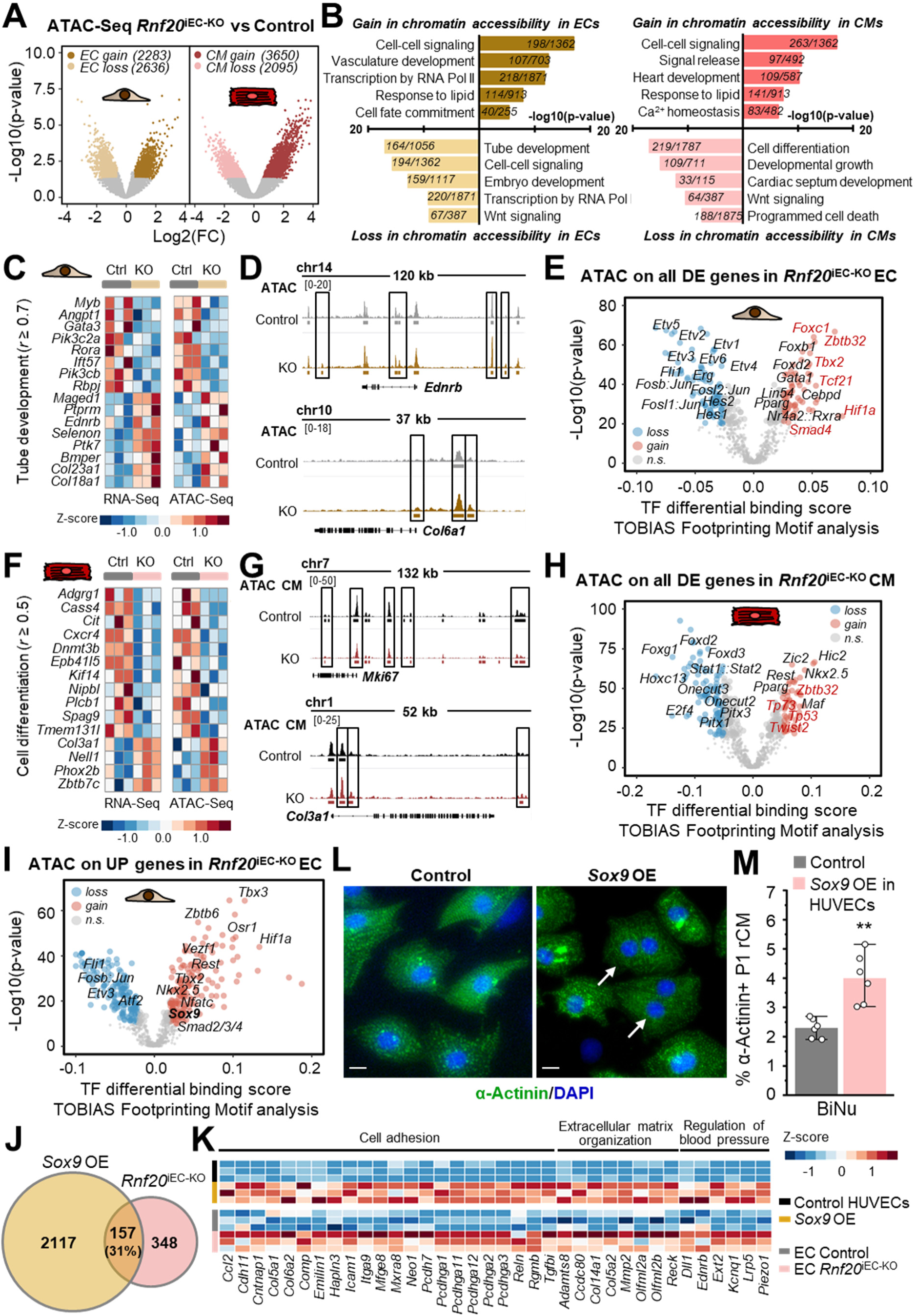
Rnf20 safeguards endothelial function by controlling TF inputs. (A) Volcano plot showing the distribution of differentially accessible chromatin regions between ECs and CMs isolated from control and Rnf20^iEC-KO^ hearts. n=3; Log2(FC)≤ -0.58, ≥0.58; p-value < 0.05. (B) Gene ontology pathway enrichment analysis of genes showing increased (top) or decreased (bottom) chromatin accessibility in ECs (left panel) and CMs (right panel) isolated from control and *Rnf20*^iEC-KO^ hearts. (C, D) Heatmap representation of genes associated to the GO: Tube development showing increased or decreased chromatin accessibility and transcriptional activity (spearman correlation factor r ≥ 0.7) in ECs isolated from control and *Rnf20*^iEC-KO^ hearts (C). Examples of genes showing increased chromatin accessibility (D). Genome tracks of ATAC-Seq reads of control and *Rnf20*^iEC-KO^ ECs are presented. (E) TOBIAS footprints of chromatin-associated proteins at ATAC-Seq peaks associated to differentially expressed (DE) genes in control and *Rnf20*^iEC-KO^ versus control ECs. (F, G) Heatmap representation of genes associated to the GO: Regulation of cell differentiation showing increased or decreased chromatin accessibility and transcriptional activity (spearman correlation factor r ≥ 0.5) in CMs isolated from control and *Rnf20*^iEC-KO^ hearts (F). Examples of genes showing decreased (top) or increased (bottom) chromatin accessibility (G). Genome tracks of ATAC-Seq reads of control and *Rnf20*^iEC-KO^ CMs are presented. (H) TOBIAS footprints of chromatin-associated proteins at ATAC-Seq peaks associated to differentially expressed (DE) genes in *Rnf20*^iEC-KO^ CMs. (I) TOBIAS footprinting analysis on all ATAC peaks associated to upregulated genes (UP) upon Rnf20 LOF in ECs. (J) Venn diagram showing the overlap of genes upregulated in *Rnf20*^iEC-KO^ versus control cardiac ECs and in HUVECs overexpressing Sox9 versus control HUVECs. For both datasets: Log2(FC) ≤ -0.58, ≥0.58; p-value < 0.05 (K) Heatmap representing gene expression levels of genes involved in cell adhesion, extracellular matrix organization and regulation of blood pressure in control and *Rnf20*^iEC-KO^ cardiac ECs and in control HUVECs and HUVECs overexpressing Sox9. (L, M) Immunostaining for α−actinin and DAPI of CMs co-cultured with HUVECs overexpressing β-gal protein or Sox9 (J) and quantification of binucleated CMs (K). Scale bars, 10 µm.

### Rnf20 inhibits aberrant signaling promoting CM binucleation, cell cycle withdrawal and irregular contractility

Since we observed a dramatic impact of Rnf20 LOF in ECs on CMs and genes involved in cell-cell signaling and ECM organization were highly deregulated in *Rnf20*^iEC-KO^, we next performed ligand-receptor analysis to pinpoint the angiocrine and juxtacrine signals from Rnf20-deficient ECs and receptors on ECs or CMs that may result in cardiac dysfunction (Figure 7A, 7B). Interestingly, we found a major increase in EC-CM signaling upon *Rnf20* LOF (Figure 7A). Among others, we identified interactions between collagens, integrins and signaling receptors together with Transforming Growth Factor Beta Induced (Tgfbi) and the tissue inhibitor of metalloproteinases-3 (Timp3), which have been shown to inhibit CM proliferation (Rhee et al., 2021), to be highly upregulated in Rnf20-deficient ECs. On the other hand, Fgf signals were decreased upon Rnf20 EC LOF (Figure 7B). Similarly, we observed increased signaling derived from CMs upon Rnf20 LOF in ECs. We next selected highly altered EC receptors, angiocrine factors and EC-derived ECM proteins to assess their role on CM proliferation and binucleation (Figure 7B). To this end, we infected HUVECs with lentiviruses carrying overexpression constructs of altered EC-derived factors as well as signaling receptors on ECs, and co-cultured them with neonatal CMs 2 days after infection (Figure 7C). Indeed, many of the selected factors blocked proliferation and/or induced CM binucleation, e.g. PCOLCE2, COL23A1, COL5A1, ICAM1, TGFBI, TIMP3 as well as TGFBR3 and PIEZO1 (Figure 7D, 7E, Figure S5A-C). In addition, we observed changes in CM beating rate and / or abnormal contractility upon overexpression of PCOLCE2, COL23A1, TIMP3, MMP14, TGFBR3 and PIEZO1 in ECs (Figure 7F, 7G, Figure S5D). Together, these data support a key function of Rnf20 in inhibiting aberrant angiocrine signaling promoting CM cell cycle withdrawal and abnormal contractility.

**Figure 7.**
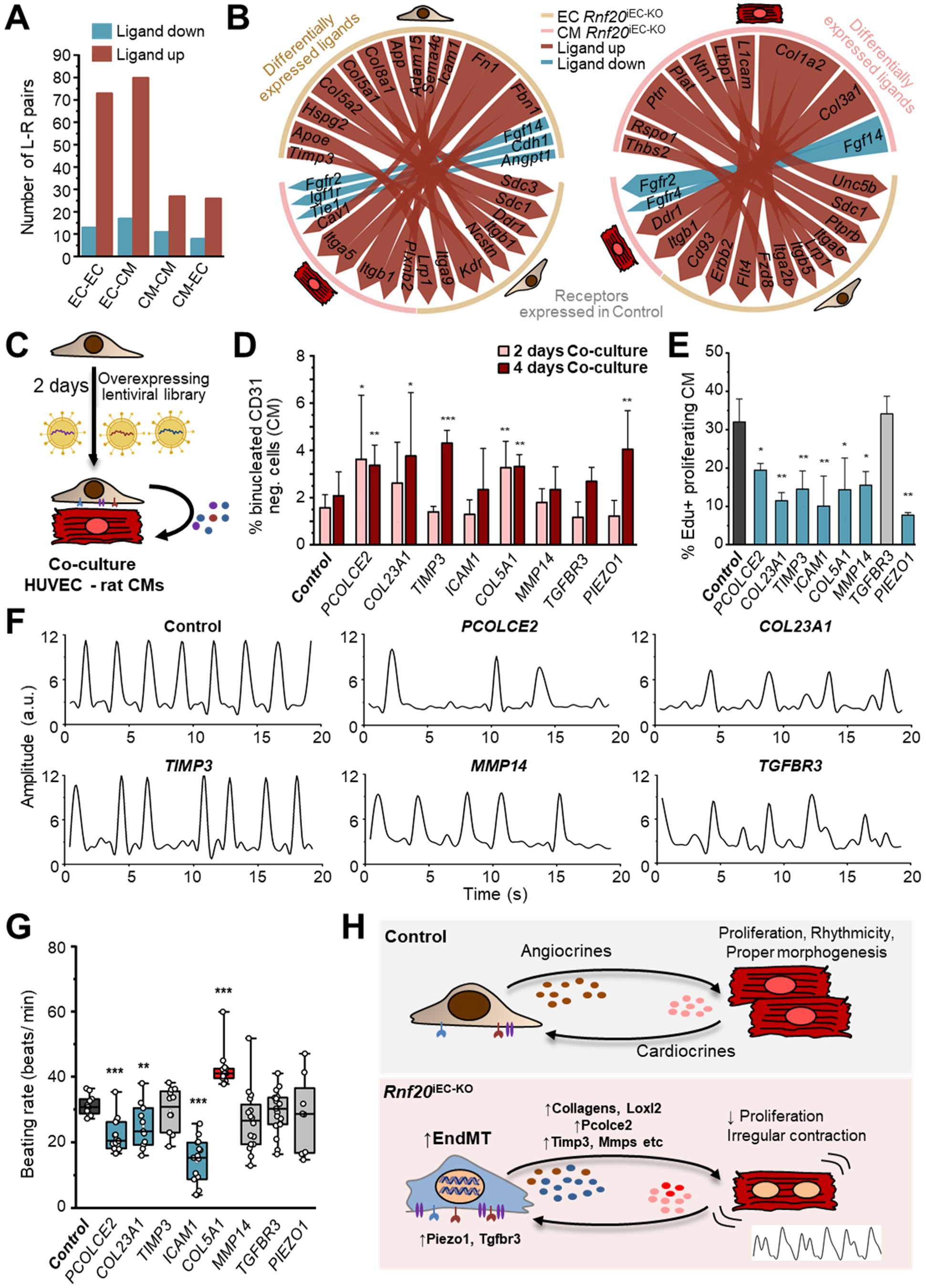
Rnf20-dependent EC-derived factors regulate CM proliferation, binucleation and contractility. (A) Bar plot of total numbers of ligand-receptor interactions between ECs and CMs including autocrine signaling in *Rnf20*^iEC-KO^ versus control (Skelly et al., 2018). (B) Circos plot of the significantly changed ligand-receptor interactions between ECs and CMs upon Rnf20 LOF in ECs. Red labeled arrow represent upregulated ligand, blue labeled downregulated ligand and arrow thickness indicates the interaction score (Skelly et al., 2018). (C) Schematic representation of the experimental design: Lentivirus particles produced by overexpression of lentiviral plasmids from genome-wide libraries in HEK293 cells were used to transduce HUVECs for 2 days. HUVECs overexpressing selected factors were then co-cultured with wild-type P1 rat CM. (D) Percentage of mononucleated and binucleated CMs determined by staining with Vybrant DyeCycle DNA dye and CD31 (to exclude HUVEC cells) and subjected to FACS analysis. P1 rat CMs were co-cultured with HUVECs overexpressing selected factors for 2 or 4 days (n=3). (E) Percentage of proliferating EdU-positive CMs determined by EdU-incorporation and FACS analysis. P1 rat CMs were co-cultured with HUVECs overexpressing selected factors for 2 days. (F) Plot of contraction amplitude and speed in spontaneously beating CMs extracted from video sequences using MYOCYTER. P1 rat CMs were co-cultured with HUVECs overexpressing selected factors for 4 days. (G) Beating rate of rat CMs co-cultured with HUVECs overexpressing selected factors. Beating rate was extracted from video sequences using MYOCYTER. (H) Model of the function of Rnf20 in ECs for proper cardiac morphogenesis and function.

## Discussion

In this study, we show that Rnf20 is a central determinant of heart development that controls EC-CM crosstalk during cardiac morphogenesis at different levels. First, Rnf20 controls extracellular matrix dynamics and cell signaling essential for second heart field development. Second, it safeguards EC identity and function by preventing EndMT. Third, it inhibits vicious angiocrine signaling and ECM deposition that cause precocious CM binucleation, cell cycle withdrawal and irregular contractility. We further identify specific factors derived from Rnf20-deficient ECs, e.g. collagens, Pcolce2, Icam1, Mmp14, Tgfbi, Tgfb3, Piezo1, etc. that modulate CM function (Figure 7H).

### Rnf20 controls extracellular matrix dynamics and cell signaling essential for second heart field development

In a protein array screen using recombinant Isl1 protein we identified Rnf20 as a putative direct interacting partner of Isl1. Deletion of Rnf20 in Isl1+ SHF progenitors resulted in major cardiovascular defects and deregulation of a multitude of common targets, supporting a key role of the Isl1-Rnf20 complex in SHF development. Interestingly, a large number of Isl1-Rnf20 target genes were linked to ECM organization and cell signaling, including genes involved in Notch signaling. Notch signaling, which plays a central role in EC function, regulates trabeculation, OFT and valve morphogenesis, and alterations in endocardial Notch signaling result in cardiac abnormalities (Del Monte-Nieto et al., 2018; Grego-Bessa et al., 2007). The Drosophila homologues of Rnf20, dBre1, was shown to be required for the expression of Notch target genes by coupling H2Bub1 to H3K4me3 and transcriptional activation in Drosophila development (Bray et al., 2005). In line with this study, we found a significant overlap between target genes of Rnf20 and Rbpj, the transcription factor mediating canonical Notch signaling. Among the common targets, particularly interesting were genes involved in ECM organization, consistent with the important role of Notch1 in the regulation of cardiac jelly dynamics (Del Monte-Nieto et al., 2018). Cardiac jelly deposition and degradation is crucial for cardiac morphogenesis. While depletion of key components of the cardiac jelly, such as versican or hyaluronan synthase-2, results in loss of trabeculation and severe cardiac chamber defects, proper degradation of the cardiac jelly by Adam and Adamts family members is also required for myocardial remodeling and valve formation (Nandadasa et al., 2014). Cardiac jelly degradation was compromised upon Rnf20 depletion, suggesting that the observed defects in myocardial growth and maturation as well as EC-CM signaling, could be due to the inability of Rnf20-deficient cells to degrade the cardiac jelly. In addition, we observed a significant decrease in Angpt1-Tie1/Tek signaling that was also shown to regulate cardiac jelly degradation (Kim et al., 2018), which could contribute to the observed excessive cardiac jelly in Rnf20 knockout hearts. Together, our data suggest an important role of the Isl1-Rnf20 complex in controlling the ECM dynamics and SHF development.

### Rnf20 maintains physiological angiocrine signaling levels that shape the structural and functional properties of the heart

Our results point to a crucial function of Rnf20 in ECs for maintaining physiological angiocrine signaling. Chromatin accessibility analysis identified genes involved in cell-cell signaling as main Rnf20 targets upregulated upon Rnf20 LOF. In the last decade it became clear that inducible transcription is mainly regulated by Pol II pausing and its release into active elongation (Hargreaves et al., 2009; Levine, 2011; Liu et al., 2015). Rnf20 and Rnf20-mediated H2Bub1 work in concert with the FACT complex to regulate elongation by Pol II (Pavri et al., 2006), but at the same time can induce transcriptional pausing by inhibiting the recruitment of TFIIS, a factor involved in the release of stalled Pol II (Shema et al., 2011). In addition, the Pol II-associated Factor 1 (PAF1), which mediates the interaction of Rnf20 with Pol II, was shown to function as a key regulator of Pol II pausing (Shema et al., 2011). Thus, Rnf20 could induce transcriptional pausing in the developing heart, thereby keeping signal-inducible transcription in check. In such a scenario, loss of Rnf20 would result in paused Pol II release and higher expression of genes involved in cell-cell communication. In line with this notion, our results revealed that Rnf20 is essential in maintaining physiological angiocrine signaling, which shapes the structural and functional properties of the heart. Indeed, inducible ablation of Rnf20 in cardiac ECs resulted in upregulation of multitude of angiocrine factors and ECM proteins coupled to major defects in CM proliferation, myocardial growth and cardiac contractility. CM dysfunction could result from altered tissue perfusion in malformed hearts. However, by using a novel cell-culture based system for co-differentiation of cardiac endothelial cells and cardiomyocytes from pluripotent stem cells, we confirmed the critical role of Rnf20 in ECs for CM proliferation and contractility. A large number of upregulated genes upon Rnf20 LOF are ECM proteins, which alter tissue stiffness and biomechanical signaling in the heart. Cardiac stiffness gradually increases during embryonic development and after birth, and maintenance of physiological levels of cardiac stiffness is detrimental for proper CM proliferation, maturation and contractility throughout life (Munch and Abdelilah-Seyfried, 2021). Systematic overexpression of selected factors using genome-wide libraries in ECs and co-culture with CMs revealed many novel molecules that regulate CM binucleation, proliferation and contractility, e.g. Pcolce2, Col5a1, Col23a1, Icam1, Mmp14, Tgfb3, Piezo1, etc., together with already known factors such as Tgfbi and Timp3 (Rhee et al., 2021). Collagens are crosslinked through lysyl oxidases (Lox, Loxl), which increases their stiffness. Together with many collagens, Loxl2 was significantly upregulated upon Rnf20 LOF, which might increase cardiac stiffness above the physiological level and thereby induce premature CM differentiation, binucleation and cell cycle withdrawal. LOXL2 is upregulated in diseased human hearts and LOXL2 inhibition protects against stress-induced cardiac dysfunction (Yang et al., 2016), suggesting that Loxl2 inhibition might be beneficial in CHD caused by Rnf20 LOF.

### Rnf20 safeguards EC identity by inhibiting EndMT

Our data revealed that Rnf20 safeguards EC identity. Rnf20-deficient EC show increased migratory capacity and upregulation of many typical markers of EndMT (Dejana et al., 2017), e.g. Fn1, Vim, Icam1, Mmp2, multiple collagens, as well as TFs driving EndMT in the heart such as Hand2, Foxc1 and Notch, suggesting that Rnf20 safeguards EC identity by preventing EndMT. TF footprint analysis at accessible chromatin regions predicted significantly increased binding of Sox9 and decreased binding of master TFs controlling EC fate, including Etv2 (Gong et al., 2022), Erg and Fli1 as well as downsteam effectors of the Notch pathway such as Hes1, required for SHF development (Rochais et al., 2009). Sox9 regulates ECM production and organization and plays an important role in EndMT in the developing heart (Akiyama et al., 2004; Hanley et al., 2008; Lincoln et al., 2007; Scharf et al., 2019). Sox9 overexpression is sufficient to activate mesenchymal genes in ECs and induce EndMT (Fuglerud et al., 2022), while silencing of Erg and Fli similarly triggers EndMT (Nagai et al., 2018). In line with increased Sox9 binding in *Rnf20*^iEC-KO^ ECs, we found a large overlap of genes involved in cell adhesion, collagen and elastic fibril organization as well as genes involved in regulation of blood pressure upon Sox9 overexpression and Rnf20 LOF. Furthermore, Sox9 overexpression in ECs resulted in increased binucleation, pointing to Sox9 as a critical mediator of Rnf20 function in the heart. Interestingly, a recent study revealed that the relative levels of Notch signaling and Sox9 control EC cell fate and plasticity in a skin wound healing experimental model (Zhao et al., 2021). We found that Rnf20 LOF reduces Notch signaling, while enhancing Sox9 binding to chromatin, suggesting that Rnf20 might be a critical regulator of this fine balance. Importantly, many Rnf20-Sox9 targets are upregulated in heart disease, where transient mesenchymal activation promotes EC migration and proliferation to regenerate the vascular network, but in the long-run might be deleterious. Thus, unraveling the mechanisms that control Rnf20 expression and function to enhance Notch and repress Sox9 activity might provide novel therapeutic avenues for cardiovascular diseases.

## Acknowledgements

We thank Dalyan Enver Devran and Alexandra Buse for technical assistance. Further, we like to thank Stefanie Uhlig, the FlowCore Mannheim, the Institute of Transfusion Medicine and Immunology and the NGS Core Facility Single Cell Services, for excellent support. We are grateful to Boyan Garvalov for careful reading of the manuscript and the constructive comments. This work was supported by the CRC 1366 (Projects A03, A06), the CRC 873 (Project A16), the CRC1550 (Project A03) funded by the DFG and the DZHK (81Z0500202), funded by BMBF.

## Author contributions

LK, RG, NT-E, OL, AZ, FT, JC, YD, YW, YR, EC, JAK, PLS, CCW performed the experiments. LK performed bioinformatic analysis, JC provided bioinformatics input. CCW, GB, TW, RO and JH provided reagents and valuable intellectual input. LK and GD designed the experiments, analyzed the data and wrote the manuscript. All authors discussed the results and commented on the manuscript.

## Methods

### Resource Availability

#### Cell lines and cell culture

HEK293T cells were purchased from ATCC (CRL-3216) and cultured in DMEM supplemented with 10% fetal bovine serum (FBS, Thermo Fisher Scientific, 10270106), 1% penicillin-streptomycin (Thermo Fisher Scientific, 15140122) and 2 mM L-glutamine (Thermo Fisher Scientific, 25030123).

HUVEC cells and cultured in Endothelial Cell Growth Medium MV 2 (PromoCell, C-22022), supplemented with 1% penicillin-streptomycin.

One day old (P1) rat CMs were isolated via Langendorff perfusion technique and cultured in PAN BIOTECH M199 media (Biolab, P04-07500), supplemented with 10% FBS.

Murine E14-Nkx2-5-EmGFP ESCs (Hsiao et al., 2008) were maintained on mitomycin (Sigma-Aldrich, M4287) treated mouse embryonic fibroblasts (MEF) in KnockOut^TM^ DMEM (Thermo Fisher Scientific, 10829018) supplemented with 10% KnockOut serum replacement (Thermo Fisher Scientific, 10828028), 2 mM L-glutamine, 0.04 mM 2-mercaptoethanol (Sigma-Aldrich, M3148), 0.1 mM non-essential amino acids (Thermo Fisher Scientific, 11140035), 1 mM sodium pyruvate (Thermo Fisher Scientific, 11360070), 4.5 mg/ml D-glucose, 1% penicillin-streptomycin and 1000 U/ml leukemia inhibitory factor (LIF ESGRO, Millipore, ESG1107).

#### Mouse lines

*Isl1* knockout mouse line (Isl1tm1(cre)Sev-C57BL/6) was a kind gift from Sylvia Evans. The Tg(Tek-cre)12Flv line was obtained from Jackson Laboratory. The Tg(Cdh5-cre/ERT2)1Rha was obtained from Prof. Ralf H. Adams. Cre deleter lines were crossed with Rnf20tm1c(EUCOMM)Wtsi mice to induce specific deletion of *Rnf20* in Isl1 positive, Tie2 positive and Cdh5 positive cells following the presented in the figures tamoxifen injection schemes. All animal experiments were performed according to the regulations issued by the Committee for Animal Rights Protection of the State of Baden-Württemberg (Regierungspraesidium Karlsruhe).

#### Isolation and culture of neonatal rat cardiomyocytes (rCM)

Hearts were collected from 1-3 days old rats, the atria removed and ventricles minced into small pieces on ice with 1x ADS buffer (pH 7.35). 50 hearts per falcon were digested with Collagenase Type II (Worthington: LS004176) and Pancreatin (Sigma P3292) and loaded to a Percoll gradient with different densities to separate into CMs, ECs and FBs. CMs (2.5×10^5^/well) were collected and seeded on plates coated with 0.5% gelatine and used for indirect co-culture or directly co-cultured with HUVECs.

#### Establishment of stable HUVEC cell lines by lentiviral infection and co-culture with rCM

HEK293T at 60-80% confluence were transfected with plasmids amplified from an overexpressing lentiviral library together with 0.975µg CMVΔR8.74 and 0.525µg VGV.G packaging plasmids using X-tremeGENE™ HP DNA Transfection Reagent (Roche, 6366236001) according to the manufacturer instructions. 48 hours after the transfection the virus-containing supernatants were used to infect HUVECs at a 30-60% confluence. Polybrene (8 mg/ml) was added 1:1000 to the infection medium. For Sox9 overexpression, HUVECs at 85-90% confluence were infected with approx. 10^6^ IFU/mL of adSox9 or adβGal particles in EC full medium for 48 hours. Afterwards, wild type rat cardiomyocytes were plated on the confluent monolayers of HUVECs at a density of 300,000 cells per well of a 6 well-plate in EC media. Contractility, binucleation and proliferation were examined at 2 or 4 days of co-culture.

#### Contraction analysis

Movies of spontaneous beating were acquired for at least 20 seconds with a Leica DMI8 microscope and a 10x or 20x objective. Contractility parameters such as beating rate and the amplitude were obtained by analysis with the ImageJ macro MYOCYTER (Grune et al., 2019). Amplitude curves were visualized by OriginPro 2020b.

#### Histology

Embryos or dissected hearts were fixed with 4% paraformaldehyde, dehydrated by increasing percentage of ethanol and stored at -20°C until paraffin embedding. Paraffin embedded samples were cut into 7 µm sections. Hematoxylin and Eosin Y staining (Sigma-Aldrich, GHS116, HT110216) as well as Alcian Blue and Nuclear Fast Red (Sigma-Aldrich, 101647, 100121) were performed according to the manufacturer’s instruction. Sections were mounted with ROTI®Histokitt (Carl Roth, 6638.1).

#### Immunofluorescence staining and imaging

For immunofluorescence staining on paraffin heart sections slides were first dewaxed and heat-induced epitope retrieval was performed by boiling for 10 minutes at 400 Watt in a pressure cooker filled with freshly prepared 10 mM sodium citrate solution with a pH of 6.0. After cooling down, the slides were blocked in a humidified chamber for 1 hour in blocking buffer consisting of 0.5% Triton X-100 and 10% FBS in PBS. The following primary antibodies were used in blocking buffer and incubated overnight at 4°C in a humidified chamber: anti-myosin heavy chain (MF20) (1:5, Dshb), anti-phospho-Histone H3 (Ser10) (1:300, Millipore, 06-570), anti-Ubiquityl-Histone H2B (Lys120) (1:100, Cell Signaling, 5546) and anti-Vimentin (1:50, Cell Signaling, 5741). After three washing steps with PBS, combinations of the corresponding secondary antibodies, conjugated to Alexa Fluor 555 or Alexa Fluor 488 (1:400, Thermo Fisher Scientific) with the conjugated primary antibody Isolectin B4 (IB4) Alexa Fluor 647 (1:40, Thermo Fisher Scientific, I32450) were added in blocking buffer and incubated for 1 hour at room temperature. The slides were then washed 3 times with PBS and stained with DAPI in blocking buffer (1:1000) for 5 minutes at room temperature, followed by 3 PBS washing steps. Sections were mounted with Mowiol 4-88 (Millipore, 475904) mounting medium.

For immunofluorescence staining on rCM cells were permeabilized for 10 minutes with 0.5% Triton X-100 in PBS. After blocking in 3% BSA in PBS for 30 min, rCM were incubated with α-Actinin (Sigma-Aldrich, A7811) antibody in 0.5% Triton X-100/ 1% BSA/ 1xPBS overnight in a humidified chamber at 4°C. After three consecutive 5 minutes washes in PBS, cells were incubated for a further 1 hour with a corresponding secondary antibody followed by DAPI staining.

Immunostaining as well as HE and Alcian Blue/Nuclear Fast Red images were acquired using Zeiss Axio Scan Z1 Digital Slide Scanner with a 10x or 20x objective and analyzed using Zen 3.5 blue software or using Leica DMi8 microscope and analyzed using Leica Application Suite X software.

#### Echocardiographic assessment of cardiac dimensions and function

Fetal echocardiography was performed with Vevo 3100 imaging system (FUJIFILM VisualSonics, Toronto, Canada) via trans-abdominal ultrasound of pregnant mice. Two-dimensional B-mode and one-dimensional M-mode tracings of E14.5 mouse embryonic hearts were recorded in a four-chambered view with a 50MHz MX700 transducer. Left- and right-ventricular diameter at end-diastole and end-systole, cardiac output, ventricular ejection fraction (EF) and fractional shortening (FS) were calculated with the integrated cardiac measurement package, where at least three consecutive cardiac cycles were used for analysis.

#### Embryonic cardiomyocyte and endothelial cell isolation

Hearts from E14.5 embryos were isolated and incubated 30 minutes at 37°C on a rotating shaker with 200 µl of digestion solution per embryo consisting out of 0.05 mg/ml DNase I (Worthington, 2139) and 1 mg/ml Collagenase I (Worthington, 4197) diluted in HBSS (Thermo Fisher Scientific, 14175053) until complete digestion. Reaction was stopped by adding DMEM medium containing 10% FBS or FACS buffer (10% FBS in PBS). For FACS sorting of samples for ATAC-Seq and RNA-Seq applications, cells were centrifuged at 400xg and 4°C for 10 minutes and resupended in 1 ml of ESC DMEM medium and incubated with 3 µl CD31 coupled Dynabeads per embryo rotating for 30 minutes at room temperature. After incubation of the cells with the Dynabeads, CD31 positive cells (Endothelial cells) were captured by a magnet. The supernatant was used for the CM fraction and the beads were washed one more time with ESC DMEM medium and twice with PBS before proceeding with ATAC-Seq and RNA isolation.

#### Flow cytometry

Cultured cells were dissociated with StemPro Accutase for 20 minutes at 37°C. Cells from E14.5 whole hearts were isolated by digestion with DNase I and Collagenase I as described before. Cells were transferred to a U-bottom-96-well plate (100 000 – 200 000 cells/well) and first stained with 50 µl/well of Fixable Viability Dye eFluor 450 (1:800, Thermo Fisher Scientific, 65-0863-14). All used antibodies were titrated with their corresponding isotypes. The following antibodies and dyes were used for extracellular staining with a FACS buffer containing 10% FBS in PBS: CD31-APC with mouse specificity (1:500, Thermo Fisher Scientific, 17-0311-82) together with its IgG2a kappa APC-Isotype control (1:500, Thermo Fisher Scientific, 17-4321-81), CD31-PE/Cyanine7 with mouse specificity (1:500, Thermo Fisher Scientific, 25-0311-82) together with its IgG2a kappa PE/Cyanine 7-Isotype control (1:500, Thermo Fisher Scientific, 25-4321-81), CD31-APC with human specificity for HUVEC experiments (1:500, Thermo Fisher Scientific, 17-0319-42) together with its IgG1 kappa APC-Isotype control (1:500, Thermo Fisher Scientific, 17-4714-82) and Vybrant DyeCycle Green Stain (1:500, Thermo Fisher Scientific, V35004). CD31 antibodies incubated for 30 minutes at 4°C or 15 minutes at room temperature. Vybrant Dye Cycle Green Stain was added for 30 minutes at 37°C followed by direct FACS measurement without further washing steps. Edu incorporation and staining with the Click-iT™ Plus EdU Alexa Fluor™ 647 Flow Cytometry Assay Kit (Thermo Fisher Scientific, C10634) was performed according to the manufacturer’s instructions with the use of the fluorescent dye from Click-iT™ EdU Cell Proliferation Kit for Imaging, Alexa Fluor™ 488 dye (Thermo Fisher Scientific, C10337). Stained cells were analyzed using an FACS Canto II Flow Cytometer (BD). Cell sorting with the FACS Aria IIu High Speed Cell Sorter (BD) was performed by the FlowCore Mannheim. Data were analyzed using FlowJo 10.7.2 software (BD).

#### 10x Genomics scRNA-Sequencing

E14.5 control and *Cdh5^CreERT2^*-Rnf20^fl/fl^ KO hearts were digested with DNase I and Collagenase I according to the isolation technique described before. Whole heart cells were FACS sorted for living cells by staining with SYTOX™ Blue Dead Cell Stain (1:2000, Thermo Fisher Scientific, S34857) and then diluted at a density of 1000 cells/µl in 0.4% BSA. 8000 cells were used for library preparation according to the Chromium Next GEM Single Cell 3’ Kit v3.1 (10xGenomics, 1000269) with the Chromium Next GEM Chip G Single Cell Kit (10xGenomics, 1000127) and sequenced on a BGISEQ-500 platform.

#### scRNA-Sequencing analysis

10x Genomics scRNA-Sequencing from control and *Cdh5^CreERT2^*-Rnf20^fl/fl^ KO hearts was analyzed using cellranger count function and refdata-gex-mm10-2020-A genome (Zheng et al., 2017). Merging of control and KO matrices to analyze expression differences in the subpopulations was done by cellranger aggr function. Resulting clusters were identified with the help of published datasets (Hill et al., 2019; Rhee et al., 2021). Clusters identified as ECs or CMs were reclustered by the use of 10x Genomics Loupe Browser to obtain a more detailed characterization these population upon Rnf20 LOF. The same procedure was done for CMs. Volcano plots of the reclustered EC populations were extracted from the Loupe Browser.

Public available scRNA-Seq (Xiong et al., 2019) gene expression lists from E8.25 embryos were downloaded from NCBI (GSE108963) and violin plots of *Rnf20* expression in NKX2.5 and Isl1 positive cells were prepared using the R package Seurat (Hao et al., 2021).

#### RNA isolation, cDNA synthesis and qPCR

RNA for RNA-Seq was isolated using the RNeasy Plus Universal Mini Kit (QIAGEN, 73404) according to the manufacturer’s instructions. For real-time PCR analysis cDNA was synthesized with the High Capacity cDNA Reverse Transcription Kit (Applied Biosystems, 4368813) and real-time qPCR was performed using either the PowerUp SYBR Green Master Mix (Applied Biosystems, A25742) or the qPCRBIO SyGreen Blue Mix Hi-ROX (PCR Biosystems, PB20.16-51). Cycle numbers were normalized to these of α-Tubulin (Tuba1a) and/or ribosomal 18S.

#### RNA-Sequencing

Right ventricle and outflow tract were dissected from E9.5 and E10.5 control and *Isl1^cre/+^*-Rnf20^-/fl^ embryos for RNA isolation and RNA-Seq. Control and *Rnf20*^iEC-KO^ ECs and CMs for sequencing applications were isolated from E14.5 embryonic hearts as described above. RNA was isolated using the RNeasy Plus Universal Mini kit (Qiagen, 73404). Quality control and library preparation were done by BGI and sequencing was performed on a BGISEQ-500 platform.

#### RNA-Sequencing data analysis

After read quality control with the MultiQC tool (Ewels et al., 2016) RNA-Seq 50 bp single-end sequencing reads were trimmed of TruSeq3-SE adapters using Trimmomatic-0.39 (Bolger et al., 2014). Trimmed reads were mapped to the mm10 reference genome using STAR (−alignIntronMin 20 −alignIntronMax 500000) (Dobin et al., 2013) and counted using the analyzeRepeats.pl function (rna mm10 -strand both -count genes –noadj) from HOMER (Heinz et al., 2010) after creating the tag directories with makeTagDirectory. Differential expression was quantified and normalized by edgeR (Robinson et al., 2010) and annotated by biomaRt (Durinck et al., 2009) using a custom script. Differentially accessible peaks were filtered by R for a p-value of 0.05 and a Log2(FoldChange) of ≥ 0.58 and ≤ -0.58. Gene ontology pathway analysis was performed using either WEB-based GEne SeT AnaLysis Toolkit (Liao et al., 2019) or DAVID Bioinformatics Resources 6.8 (Huang da et al., 2009). The PCA plots were obtained using prcomp function of R package ggfortify (Yuan Tang, 2016) and visualization by ggplot2 (Wickham, 2016). Volcano plots were designed using EnhancedVolcano R package (Blighe K, 2020). Gene Set Enrichment Analysis (GSEA) was performed by the use of the R packages clusterProfiler (Wu et al., 2021), enrichplot (Yu, 2021) and ggplot2. All heatmaps were prepared by the R packages pheatmap (Kolde, 2019) and RColorBrewer (Neuwirth, 2014) and represent the row-based Z-scores calculated from normalized counts. Ligand-Receptor analysis was based on the database from Skelly et al. (2018). In detail, differentially regulated ligands upon *Rnf20* depletion in the different cell types were paired with expressed receptors. Further, an interaction score based on the Log2FC of the differentially expressed ligand and on the expression levels of both the ligand and receptor was calculated to sort the ligand-receptor pairs by relevance. Interactions of ligand-receptor pairs were visualized with the help of the R package circlize (Gu et al., 2014). All data were analyzed with the same parameters.

#### ATAC-Sequencing and RNA harvesting for RNA-Sequencing of E14.5 Rnf20^iEC-KO^ cells

ECs and CMs from *Cdh5^CreERT2^*-Rnf20^fl/fl^ E14.5 embryo hearts were processed using a modified protocol from Corces et al. (2017). In brief, 30 000 cells of CMs and 10 000 cells of ECs were centrifuged at 500xg for 5 minutes at 4°C and washed with PBS. The cell pellet was resuspended in 50 µL cold lysis buffer (10 mM Tris-HCl pH 7.4, 10 mM NaCl, 3 mM MgCl_2_, 0.1% Igepal CA-630, Sigma-Aldrich, I8896) and incubated for 2 minutes on ice, followed by stopping the reaction by dilution with 300 µl ATAC resuspension buffer (10 mM Tris-HCl pH 7.4, 10 mM NaCl, 3 mM MgCl_2_) and 10 minutes centrifugation step (500g, 4°C). After centrifugation supernatant was used for RNA cytosolic fraction by adding to 1 ml of TRIZOL and isolated nuclei of CMs were incubated in 30 µl and of EC in 10 µl transposase mixture with 5% Nextera Tagment DNA Enzyme TDE (Illumina #15027916) in transposition buffer (20 mM Tris-HCl pH 7.6, 10 mM MgCl_2_, 20% Dimethylformamide) under shaking for 30 minutes at 37 °C. Immediately following the transposition reaction, the mixture was purified using ChIP DNA Clean and Concentrator Kit (Zymo D5205). Library amplification together with indexing was performed by a 5 cycle preamplification step, a SYBR green qPCR side reaction with 1/10 of the preamplified solution to determine the extra cycle number and a final PCR with individually adapted cycle numbers to avoid duplicates and ensure similar final concentrations using NEBNext High-Fidelity 2x PCR Master Mix (New England Biolabs, M0541S) and Nextera Index Kit (Illumina, 15055290). Magnetic bead purification with two-sided size selection using undiluted Agencourt AMPure XP Beads (Beckman Coulter, A63881) ensured library sizes between 150 and 1000 bp, detected by Bioanalyzer High Sensitivity DNA analysis (Agilent, 5067-4626). Libraries were mixed in equimolar ratios and sequencing was performed on a BGISEQ-500 platform.

#### ATAC-Sequencing data analysis

ATAC-Seq paired-end sequencing reads were trimmed of adapters using Trimmomatic-0.39 and then mapped to mm10 mouse genome with Bowtie2 (Langmead and Salzberg, 2012). Following removal of unmapped reads by SAMtools (-F 1804 -f 2) (Li et al., 2009), PCR artefacts were excluded by the MarkDuplicates.jar from Picard-tools-1.119 (Tools, 2015). Read quality was controlled by the MultiQC tool (Ewels et al., 2016). BamCoverage function of deepTools (Ramírez et al., 2014) with normalization to the reads per genomic content was used to generate the bigwig files for visualization of ATAC-Seq reads in the genome browser IGV (-bs 20 --smoothLength 40 --normalizeUsing RPGC --effectiveGenomeSize 2150570000 -e). Therefore, bam files of each 3 replicates were merged by BamTools (Barnett et al., 2011) and processing was performed like described. Peak calling was applied from merged bam-files with MACS14 (-p 1e-3 -g 1.87e9 --nomodel --shiftsize 100) (Zhang et al., 2008). Called peaks from merged files and individual bam files were further used to quantify and normalize counts as well as calculating differentially accessible chromatin regions upon the different treatments using the R package DiffBind (Stark and Brown) with an edgeR based TMM normalization using read counts and effective library size and an edgeR method to perform differential binding affinity analysis. Peaks were annotated by a custom R script combining ChIPseeker (Yu et al., 2015) and rtracklayer packages (Lawrence et al., 2009). Promoter regions were defined as ±3 kb around the gene transcription start site. Differentially accessible peaks were filtered by R for a p-value of 0.05 and a Log2(FoldChange) of ≥ 0.58 and ≤ -0.58.

Principle component analysis (PCA) and heatmap of 10 000 most significant peaks were generated by the DiffBind package. Volcano plots are based on the EnhancedVolcano package (Blighe K, 2020). Gene ontology pathway analysis was performed using WEB-based GEne SeT AnaLysis Toolkit (Liao et al., 2019). Average ATAC-Seq tag intensities in the ATAC-Seq normalized mapped reads were calculated using ngs.plot.r (Shen et al., 2014) by filtering the ATAC-Seq peaks by the genes of interest. TOBIAS Footprinting motif analysis (Bentsen et al., 2020b) of the filtered peaks was performed with the TOBIAS BINDetect function using the merged bigwig files of the treatments and the JASPAR 2020 open-access database of curated, non-redundant transcription factor binding profiles (Fornes et al., 2019). Spearman correlation analysis of the normalized counts from all individual replicates of ATAC-Seq and RNA-Seq was based on the calculation of the spearman correlation factor r using the cor.test function in R. Visualization of the correlating genes filtered by r ≥ 0.5 and ≤ -0.5 was done by the R package ggpubr (Kassambara, 2020).

#### ChIP-Sequencing data analysis of public data sets

RNA Polymerase II (Pol II) ChIP-Seq from E14.5 hearts (Shen et al., 2012) was analyzed based on the ATAC-Seq data processing. Peak calling was applied from merged bam-files with MACS14 (-c $Input -p 1e-2 -g 1.87e9 --nomodel --shiftsize 100) considering the ChIP input as a background. Pol II ChIP-Seq peaks were filtered by differentially expressed genes upon Isl1-mediated *Rnf20* deletion as well as non-changed genes (NC) and pausing index was calculated by the ratio of -50bp to +300bp Pol II signal divided by the Pol II signal within +300bp to +2kb for these peaks. The pausing index was then classified into high (> 3^rd^ quartile), medium (between 3^rd^ quartile and median) and low (< median).

### Quantification and statistical analysis

All the statistical analyses were performed using OriginPro 2020b and R programming software. All experiments were performed at least three independent times and the respective data were used for statistical analyses. Differences between groups were assessed using an unpaired two-tailed Student’s t-test. Error bars represent mean ± SD. P-values represent statistical significance *p<0.05 **p<0.01 ***p<0.001.

## Supplemental information

**Figure S1.**
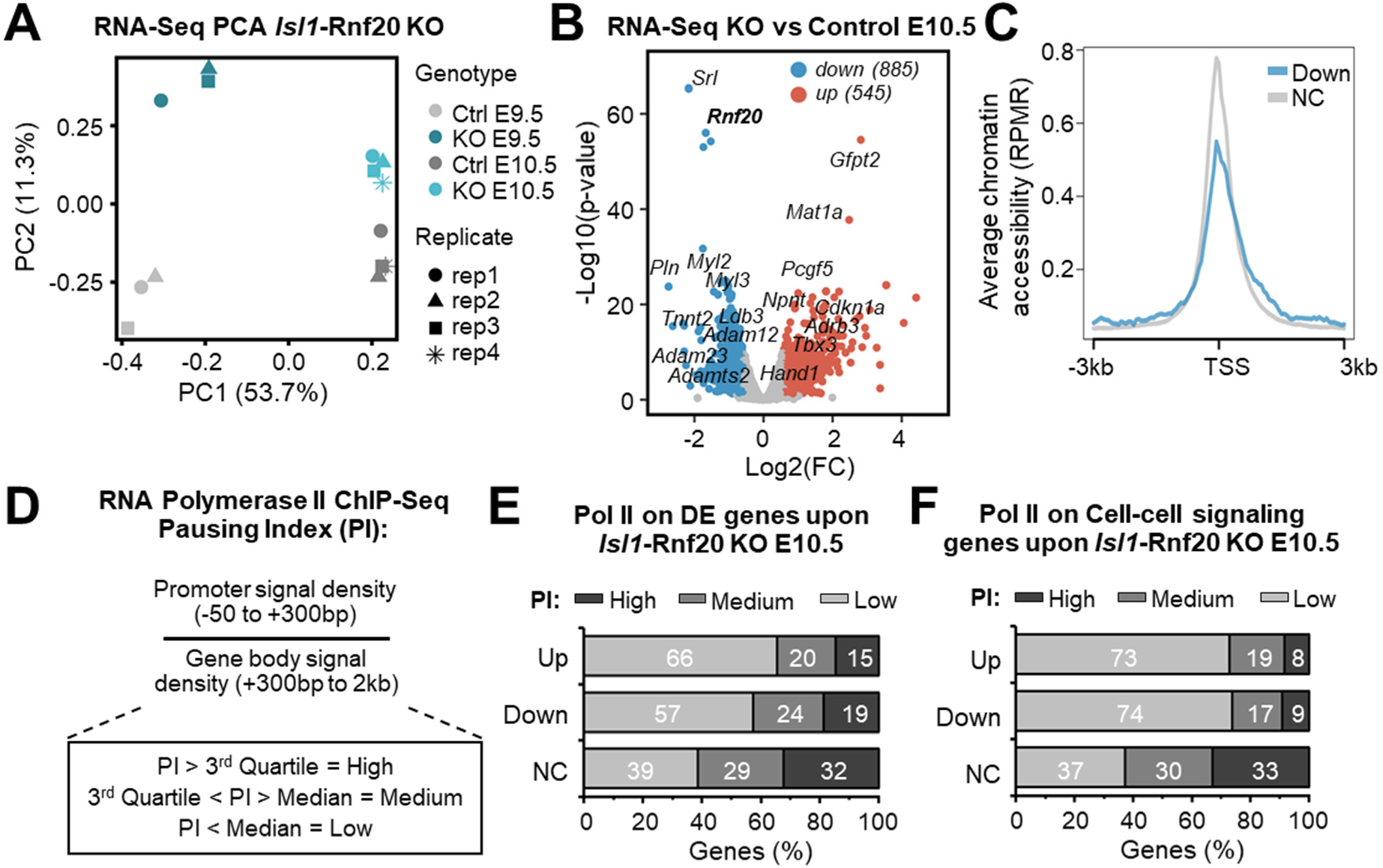
Role of Rnf20 in SHF development, Related to Figure 2. ***(A)*** Principal component analysis (PCA) of gene expression variation in dissected OFT and RV of E9.5 (n=3) and E10.5 (n=4) control and *Isl1^cre/+-^*-Rnf20^-/fl^ embryos. ***(B)*** Volcano plot showing the distribution of differentially expressed genes in dissected E10.5 OFT and RV of Isl1^cre/+^Rnf20^-/fl^ versus control OFT and RV (n=3; Log2(FC) ≤ -0.58, ≥0.58; p-value < 0.05). Representative up- (red) and downregulated (blue) genes are indicated. ***(C)*** Average ATAC-Seq tag intensities in pharyngeal mesoderm/hearts of wild-type and Isl1-/-embryos at genes downregulated upon Isl1 loss of function in E10.5 (log2FC ≤ -1, p-value < 0.05). ***(D)*** Schematic representation of Pol II average profiles and the method used for defining the Pol II pausing index. Pausing index was calculated by the ratio of -50bp to +300bp Pol II signal divided by the Pol II signal within +300bp to +2kb. ***(E)*** Pol II pausing index of upregulated, downregulated and non-changed genes in dissected OFT and RV of E10.5 control and *Isl1^cre/+-^*-Rnf20^-/fl^ embryos, showing that differentially regulated genes are associated with higher elongation rates. Pol II ChIP-Seq data from E14.5 hearts was used (GSE29218) ***(F)*** Pol II pausing index of upregulated, downregulated and non-changed cell-cell-signaling genes (GO:0007267).

**Figure S2.**
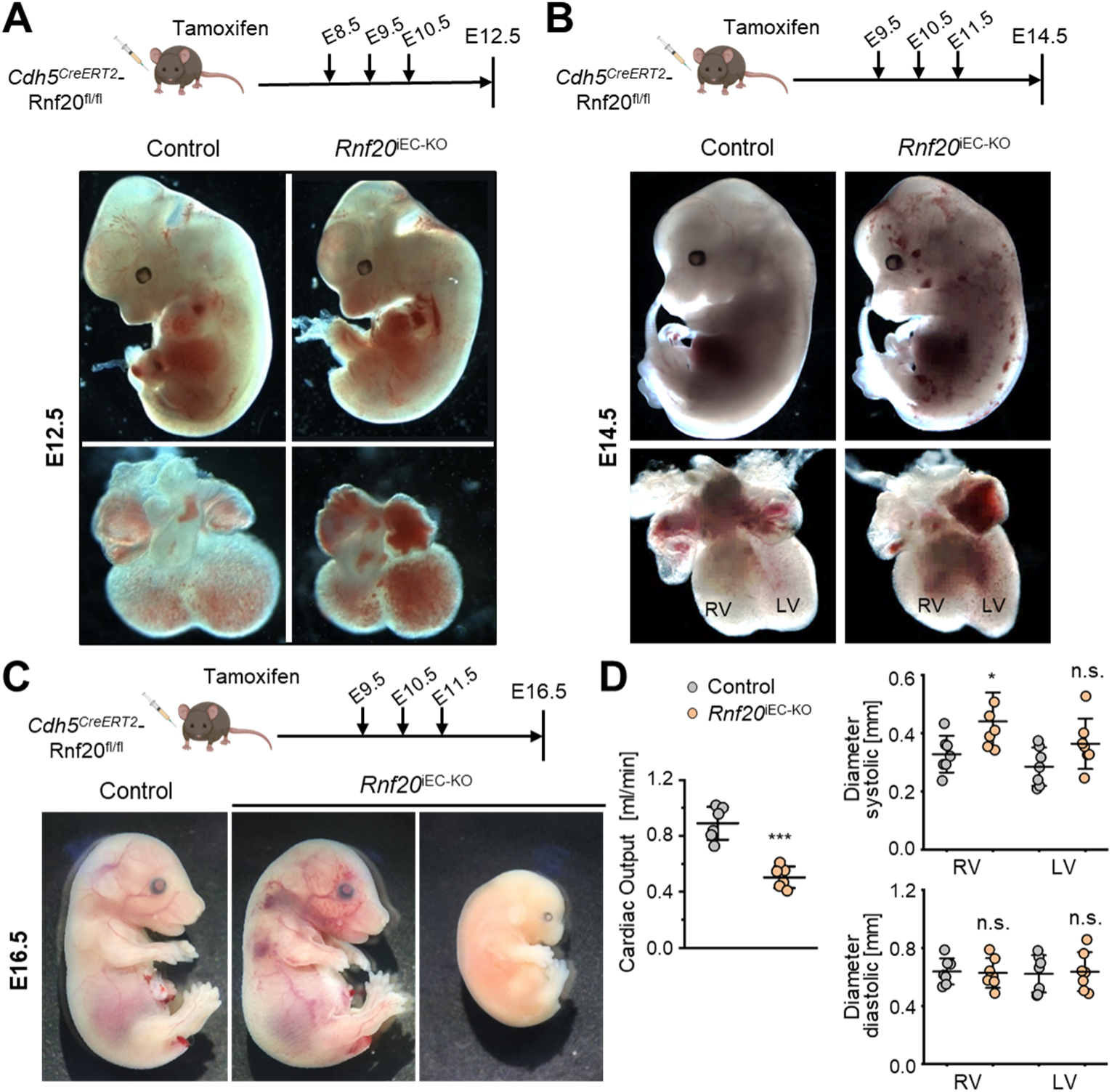
Cardiac Phenotype of control and *Rnf20*^iEC-KO^ Mice, Related to Figure 3. ***(A)*** Experimental regimen to induce Rnf20 ablation (top). First tamoxifen administration in this experiment was conducted at E8.5. Gross appearance of control (*Cdh5^CreERT2^*neg-Rnf20^fl/fl^) and *Rnf20*^iEC-KO^ (*Cdh5*^CreERT2^pos-Rnf20^fl/fl^) embryos (middle panels) and dissected hearts (bottom panels) at E12.5. ***(B)*** Experimental regimen to induce Rnf20 ablation (top). First tamoxifen administration for this experiment and all the following experiments was conducted at E9.5. Gross appearance of control (*Cdh5^CreERT2^*neg-Rnf20^fl/fl^) and *Rnf20*^iEC-KO^ (*Cdh5*^CreERT2^pos-Rnf20^fl/fl^) embryos (middle panels) and dissected hearts (bottom panels) at E14.5. ***(C)*** Experimental regimen to induce Rnf20 ablation (top). Gross appearance of control (*Cdh5^CreERT2^*neg-Rnf20^fl/fl^) and *Rnf20*^iEC-KO^ (*Cdh5*^CreERT2^pos-Rnf20^fl/fl^) embryos (bottom) at E16.5. ***(D)*** Cardiac output assessed by echocardiography of control and *Rnf20*^iEC-KO^ hearts (n=6).

**Figure S3.**
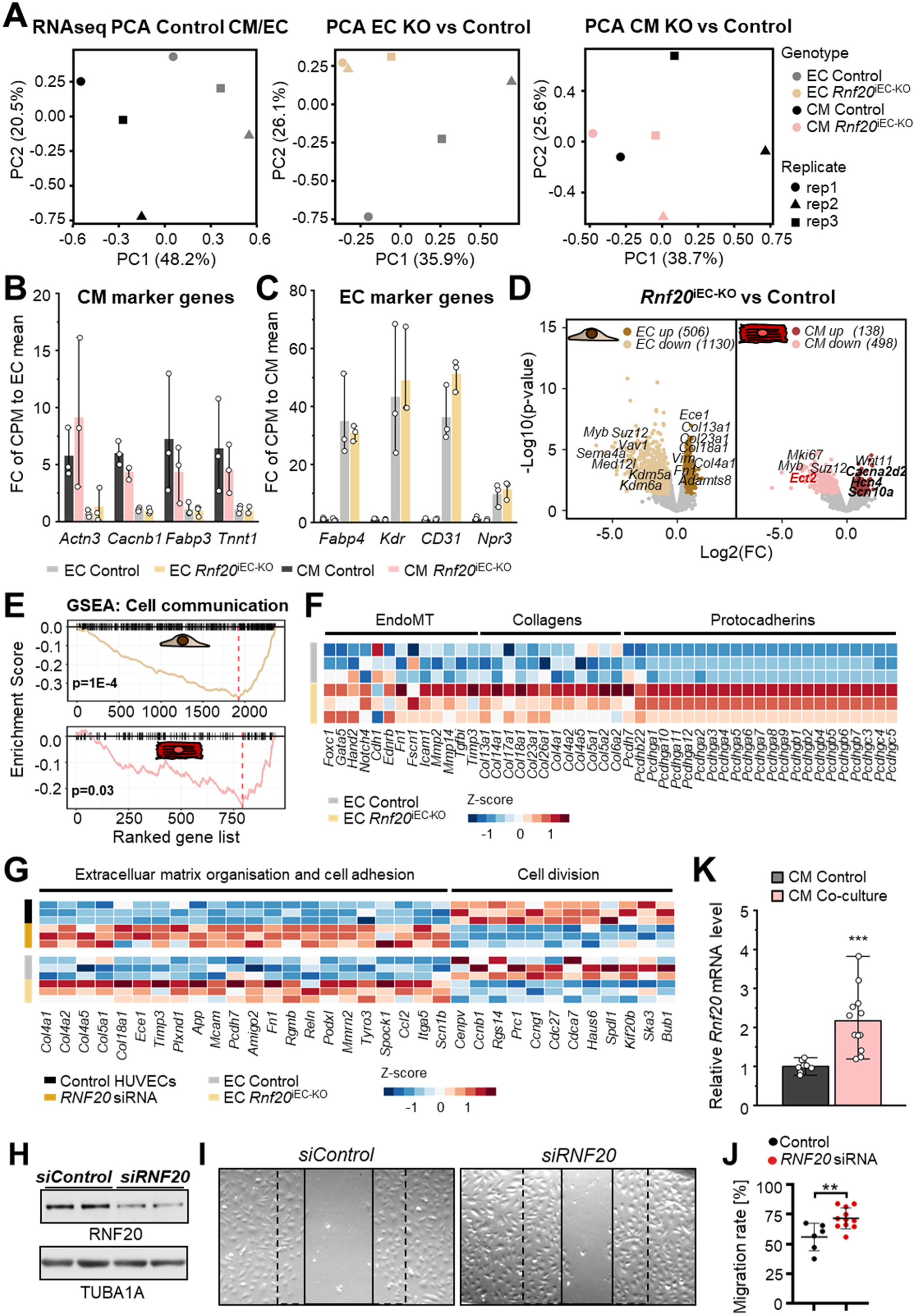
Transcriptional alterations in ECs and CMs isolated from control and Rnf20iECKO hearts, Related to Figure 5. ***(A)*** Principal component analysis (PCA) of gene expression variation of 1000 most significant genes in ECs and CMs isolated from control embryos (left), showing a clear separation between the different cell populations. PCA of gene expression variation of 1000 most significant genes in ECs (middle) and CMs (right) isolated from control and *Rnf20*^iEC-KO^ hearts. ***(B)*** Fold-change (FC) of normalized CPM expression of CM marker genes in ECs and CMs isolated from control and *Rnf20*^iEC-KO^ hearts. ***(C)*** FC of normalized CPM expression of EC marker genes in ECs and CMs isolated from control and *Rnf20*^iEC-KO^ hearts. ***(D)*** Volcano plots showing the distribution of differentially expressed genes between *Cdh5^pos^*-Rnf20^fl/fl^ and control ECs (left, brown) and CM (right, red). Representative up- (dark color) and downregulated (light) genes are indicated. n=3; Log2(FC) ≤ -0.58, ≥ 0.58; p-value < 0.05. ***(E)*** Gene set enrichment analysis of the pathway *cell communication* in downregulated genes in EC (top) and CM (bottom) upon endothelial deletion of *Rnf20*. ***(F)*** Heatmap representing gene expression levels of collagens, protocadherins and factors involved in EndMT in control and *Rnf20*^iEC-KO^ ECs. ***(G)*** Heatmap representing gene expression levels of genes involved in extracellular matrix organization and cell adhesion and cell division in control and *Rnf20*^iEC-KO^ cardiac ECs and HUVECs transfected either with control siRNA or siRNA against *Rnf20*. ***(H)*** Western blot analysis for RNF20 in HUVECs transfected with siRNA against RNF20 versus control siRNA transfected cells for the analysis in ***(I, J)***. ***(I, J)*** Scratch wound healing assay with control and RNF20 knockdown HUVECs ***(I)*** and quantification of the migration rate of HUVECs transfected with control siRNA or siRNA against *RNF20 **(J)***. ***(K)*** Relative *Rnf20* expression levels in sorted ESC-derived CMs cultured alone or together with ESC-derived endocardial ECs, showing increased Rnf20 expression in CMs upon co-culture with ECs.

**Figure S4.**
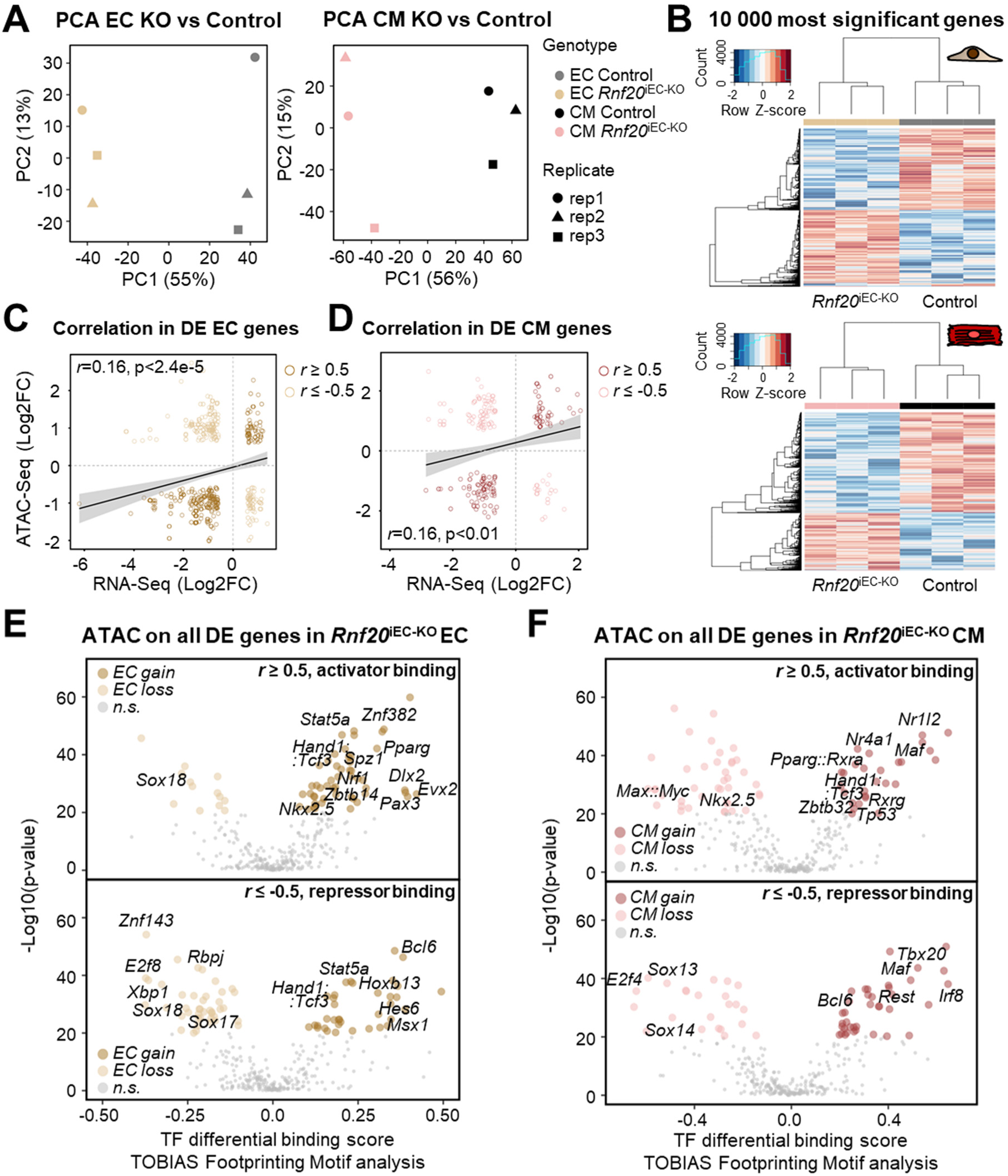
Chromatin accessibility changes in ECs and CMs upon Rnf20 LOF in ECs, Related to Figure 6. ***(A)*** Principal component analysis (PCA) of genome-wide chromatin accessibility variation in control and *Rnf20*^iEC-KO^ ECs (left panel) and CMs (right panel). ***(B*)** Heatmap of 10000 most significantly altered ATAC-Seq peaks between *Rnf20*^iEC-KO^ and control ECs (top) and CMs (bottom). ***(C)*** Scatter plot showing the spearman correlation between chromatin accessibility and gene expression changes (Log2FC) in *Rnf20*^iEC-KO^ vs control ECs for all annotated differentially expressed genes. Pre-filtering of p < 0.05 and Log2(FC) ≤ -0.58 and ≥ 0.58 was applied. Dark brown dots indicate a positive whereas light brown dots indicate a negative correlation. ***(D)*** Scatter plot showing the spearman correlation between chromatin accessibility and gene expression changes (Log2(FC)) in *Rnf20*^iEC-KO^ vs control CMs for all annotated differentially changed genes. Pre-filtering of p < 0.05 and Log2(FC) ≤ -0.58 and ≥ 0.58 was applied. Dark red dots indicate a positive whereas light red dots indicate a negative correlation. ***(E)*** TOBIAS footprints of chromatin-associated proteins at ATAC-Seq peaks on genes that show positive (top) or negative (bottom) correlation between chromatin accessibility with transcriptional activity in ECs, potentially acting as transcriptional activators or repressors, respectively. ***(F)*** TOBIAS footprints of chromatin-associated proteins at ATAC-Seq peaks on genes that show positive (top) or negative (bottom) correlation between chromatin accessibility with transcriptional activity in CMs, potentially acting as transcriptional activators or repressors, respectively.

**Figure S5.**
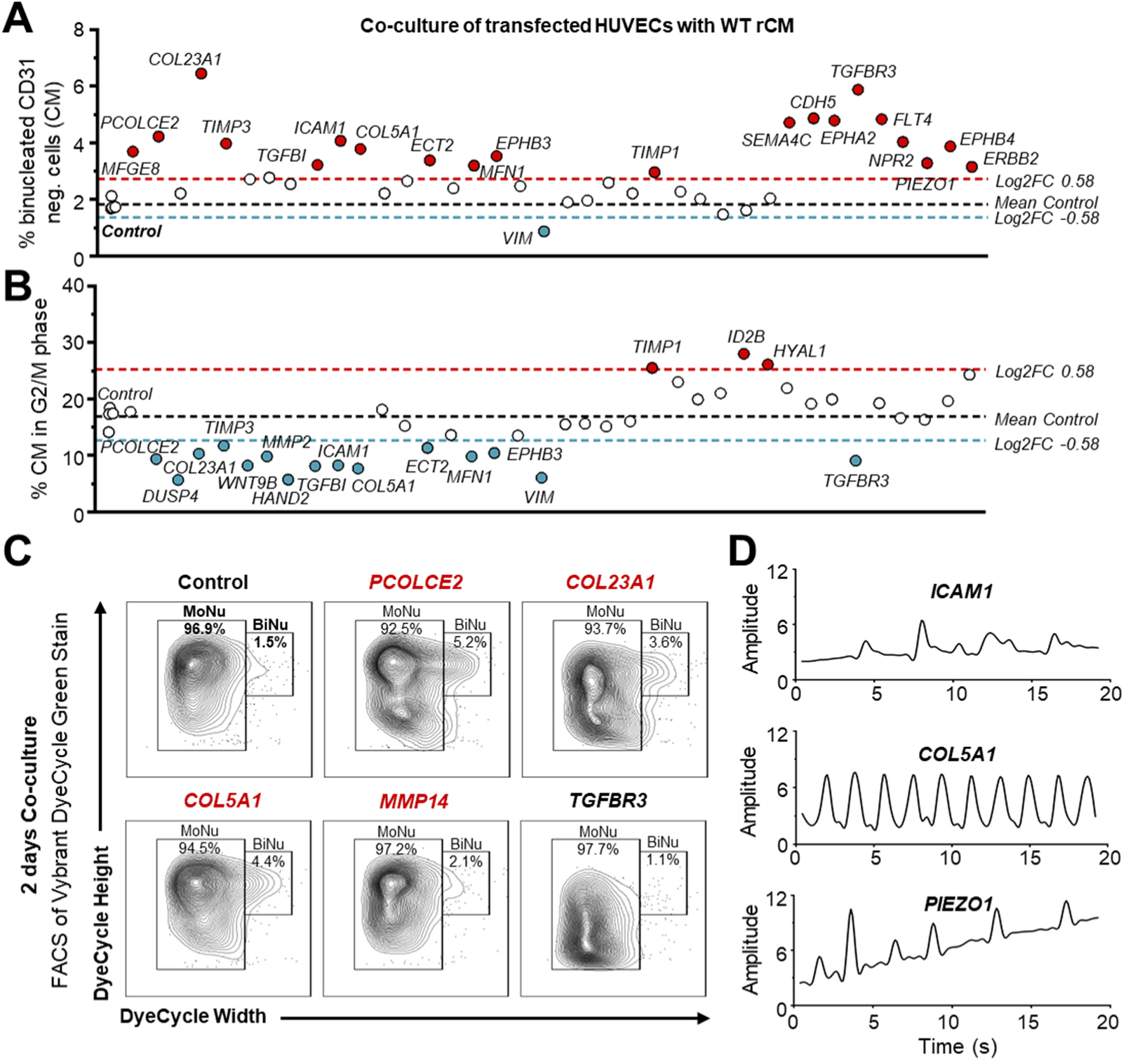
Effect of angiocrine factors, receptors on endothelial cells and extracellular matrix proteins on CM behavior, Related to Figure 7. ***(A)*** Screening: Percentage of binucleated CMs as determined by the FACS analysis with the Vybrant DyeCycle DNA dye and CD31 (to subtract HUVECs). P1 rat cardiomyocytes were co-cultured with HUVECs overexpressing selected factors. ***(B)*** Screening: Percentage of CMs in G2/M as determined by the FACS analysis with the Vybrant DyeCycle DNA dye and CD31 (to subtract HUVECs). P1 rat cardiomyocytes were co-cultured with HUVECs overexpressing selected factors. ***(C)*** Representative FACS analysis of co-cultures stained with the Vybrant DyeCycle DNA dye and CD31 (to subtract HUVECs). P1 rat cardiomyocytes were co-cultured with HUVECs overexpressing selected factors for 2 days. ***(D)*** Plot of contraction amplitude and speed in spontaneously beating CMs from video sequences using MYOCYTER. P1 rat cardiomyocytes were co-cultured with HUVECs overexpressing selected factors for 4 days.

## References

Akiyama, H., Chaboissier, M.C., Behringer, R.R., Rowitch, D.H., Schedl, A., Epstein, J.A., and de Crombrugghe, B. (2004). Essential role of Sox9 in the pathway that controls formation of cardiac valves and septa. Proc Natl Acad Sci U S A 101, 6502–6507.

Barnett, D.W., Garrison, E.K., Quinlan, A.R., Stromberg, M.P., and Marth, G.T. (2011). BamTools: a C++ API and toolkit for analyzing and managing BAM files. Bioinformatics 27, 1691–1692.

Bentsen, M., Goymann, P., Schultheis, H., Klee, K., Petrova, A., Wiegandt, R., Fust, A., Preussner, J., Kuenne, C., Braun, T., et al. (2020a). ATAC-seq footprinting unravels kinetics of transcription factor binding during zygotic genome activation. Nat Commun 11, 4267.

Bentsen, M., Goymann, P., Schultheis, H., Klee, K., Petrova, A., Wiegandt, R., Fust, A., Preussner, J., Kuenne, C., Braun, T., et al. (2020b). ATAC-seq footprinting unravels kinetics of transcription factor binding during zygotic genome activation. Nat Commun 11, 4267.

Blighe K, R.S., Lewis M (2020). EnhancedVolcano: Publication-ready volcano plots with enhanced colouring and labeling. https://githubcom/kevinblighe/EnhancedVolcano.

Bolger, A.M., Lohse, M., and Usadel, B. (2014). Trimmomatic: a flexible trimmer for Illumina sequence data. Bioinformatics 30, 2114–2120.

Bray, S., Musisi, H., and Bienz, M. (2005). Bre1 is required for Notch signaling and histone modification. Dev Cell 8, 279–286.

Buenrostro, J.D., Giresi, P.G., Zaba, L.C., Chang, H.Y., and Greenleaf, W.J. (2013). Transposition of native chromatin for fast and sensitive epigenomic profiling of open chromatin, DNA-binding proteins and nucleosome position. Nat Methods 10, 1213–1218.

Cai, C.L., Liang, X., Shi, Y., Chu, P.H., Pfaff, S.L., Chen, J., and Evans, S. (2003). Isl1 identifies a cardiac progenitor population that proliferates prior to differentiation and contributes a majority of cells to the heart. Dev Cell 5, 877–889.

Caputo, L., Witzel, H.R., Kolovos, P., Cheedipudi, S., Looso, M., Mylona, A., van, I.W.F., Laugwitz, K.L., Evans, S.M., Braun, T., et al. (2015). The Isl1/Ldb1 Complex Orchestrates Genome-wide Chromatin Organization to Instruct Differentiation of Multipotent Cardiac Progenitors. Cell Stem Cell 17, 287–299.

Chen, F.X., Woodfin, A.R., Gardini, A., Rickels, R.A., Marshall, S.A., Smith, E.R., Shiekhattar, R., and Shilatifard, A. (2015). PAF1, a Molecular Regulator of Promoter-Proximal Pausing by RNA Polymerase II. Cell 162, 1003–1015.

Claxton, S., Kostourou, V., Jadeja, S., Chambon, P., Hodivala-Dilke, K., and Fruttiger, M. (2008). Efficient, inducible Cre-recombinase activation in vascular endothelium. Genesis 46, 74–80.

Corces, M.R., Trevino, A.E., Hamilton, E.G., Greenside, P.G., Sinnott-Armstrong, N.A., Vesuna, S., Satpathy, A.T., Rubin, A.J., Montine, K.S., Wu, B., et al. (2017). An improved ATAC-seq protocol reduces background and enables interrogation of frozen tissues. Nat Methods 14, 959–962.

Dejana, E., Hirschi, K.K., and Simons, M. (2017). The molecular basis of endothelial cell plasticity. Nat Commun 8, 14361.

Del Monte-Nieto, G., Ramialison, M., Adam, A.A.S., Wu, B., Aharonov, A., D’Uva, G., Bourke, L.M., Pitulescu, M.E., Chen, H., de la Pompa, J.L., et al. (2018). Control of cardiac jelly dynamics by NOTCH1 and NRG1 defines the building plan for trabeculation. Nature 557, 439–445.

Dobin, A., Davis, C.A., Schlesinger, F., Drenkow, J., Zaleski, C., Jha, S., Batut, P., Chaisson, M., and Gingeras, T.R. (2013). STAR: ultrafast universal RNA-seq aligner. Bioinformatics 29, 15–21.

Durinck, S., Spellman, P.T., Birney, E., and Huber, W. (2009). Mapping identifiers for the integration of genomic datasets with the R/Bioconductor package biomaRt. Nat Protoc 4, 1184–1191.

Elsherbiny, A., and Dobreva, G. (2021). Epigenetic memory of cell fate commitment. Curr Opin Cell Biol 69, 80–87.

Ewels, P., Magnusson, M., Lundin, S., and Käller, M. (2016). MultiQC: summarize analysis results for multiple tools and samples in a single report. Bioinformatics 32, 3047–3048.

Feulner, L., van Vliet, P.P., Puceat, M., and Andelfinger, G. (2022). Endocardial Regulation of Cardiac Development. J Cardiovasc Dev Dis 9.

Fornes, O., Castro-Mondragon, J.A., Khan, A., van der Lee, R., Zhang, X., Richmond, P.A., Modi, B.P., Correard, S., Gheorghe, M., Baranašić, D., et al. (2019). JASPAR 2020: update of the open-access database of transcription factor binding profiles. Nucleic Acids Research 48, D87–D92.

Fuchs, G., and Oren, M. (2014). Writing and reading H2B monoubiquitylation. Biochim Biophys Acta 1839, 694–701.

Fuglerud, B.M., Drissler, S., Lotto, J., Stephan, T.L., Thakur, A., Cullum, R., and Hoodless, P.A. (2022). SOX9 reprograms endothelial cells by altering the chromatin landscape. Nucleic Acids Res.

Gao, R., Liang, X., Cheedipudi, S., Cordero, J., Jiang, X., Zhang, Q., Caputo, L., Gunther, S., Kuenne, C., Ren, Y., et al. (2019). Pioneering function of Isl1 in the epigenetic control of cardiomyocyte cell fate. Cell Res 29, 486–501.

Gong, W., Das, S., Sierra-Pagan, J.E., Skie, E., Dsouza, N., Larson, T.A., Garry, M.G., Luzete-Monteiro, E., Zaret, K.S., and Garry, D.J. (2022). ETV2 functions as a pioneer factor to regulate and reprogram the endothelial lineage. Nat Cell Biol 24, 672–684.

Grego-Bessa, J., Luna-Zurita, L., del Monte, G., Bolos, V., Melgar, P., Arandilla, A., Garratt, A.N., Zang, H., Mukouyama, Y.S., Chen, H., et al. (2007). Notch signaling is essential for ventricular chamber development. Dev Cell 12, 415–429.

Grune, T., Ott, C., Häseli, S., Höhn, A., and Jung, T. (2019). The “MYOCYTER” – Convert cellular and cardiac contractions into numbers with ImageJ. Scientific Reports 9, 15112.

Gu, Z., Gu, L., Eils, R., Schlesner, M., and Brors, B. (2014). circlize Implements and enhances circular visualization in R. Bioinformatics 30, 2811–2812.

Hanley, K.P., Oakley, F., Sugden, S., Wilson, D.I., Mann, D.A., and Hanley, N.A. (2008). Ectopic SOX9 mediates extracellular matrix deposition characteristic of organ fibrosis. J Biol Chem 283, 14063–14071.

Hao, Y., Hao, S., Andersen-Nissen, E., Mauck, W.M., Zheng, S., Butler, A., Lee, M.J., Wilk, A.J., Darby, C., Zager, M., et al. (2021). Integrated analysis of multimodal single-cell data. Cell 184, 3573–3587.e3529.

Hargreaves, D.C., Horng, T., and Medzhitov, R. (2009). Control of inducible gene expression by signal-dependent transcriptional elongation. Cell 138, 129–145.

Heinz, S., Benner, C., Spann, N., Bertolino, E., Lin, Y.C., Laslo, P., Cheng, J.X., Murre, C., Singh, H., and Glass, C.K. (2010). Simple combinations of lineage-determining transcription factors prime cis-regulatory elements required for macrophage and B cell identities. Mol Cell 38, 576–589.

Hill, M.C., Kadow, Z.A., Li, L., Tran, T.T., Wythe, J.D., and Martin, J.F. (2019). A cellular atlas of Pitx2-dependent cardiac development. Development 146.

Hsiao, E.C., Yoshinaga, Y., Nguyen, T.D., Musone, S.L., Kim, J.E., Swinton, P., Espineda, I., Manalac, C., deJong, P.J., and Conklin, B.R. (2008). Marking embryonic stem cells with a 2A self-cleaving peptide: a NKX2-5 emerald GFP BAC reporter. PLoS One 3, e2532.

Huang da, W., Sherman, B.T., and Lempicki, R.A. (2009). Systematic and integrative analysis of large gene lists using DAVID bioinformatics resources. Nat Protoc 4, 44–57.

Kassambara, A. (2020). ggpubr: ’ggplot2’ Based Publication Ready Plots. https://CRANR-projectorg/package=ggpubr.

Kim, H., Wang, M., and Paik, D.T. (2021). Endothelial-Myocardial Angiocrine Signaling in Heart Development. Front Cell Dev Biol 9, 697130.

Kim, J., Guermah, M., McGinty, R.K., Lee, J.S., Tang, Z., Milne, T.A., Shilatifard, A., Muir, T.W., and Roeder, R.G. (2009). RAD6-Mediated transcription-coupled H2B ubiquitylation directly stimulates H3K4 methylation in human cells. Cell 137, 459–471.

Kim, J., Hake, S.B., and Roeder, R.G. (2005). The human homolog of yeast BRE1 functions as a transcriptional coactivator through direct activator interactions. Mol Cell 20, 759–770.

Kim, K.H., Nakaoka, Y., Augustin, H.G., and Koh, G.Y. (2018). Myocardial Angiopoietin-1 Controls Atrial Chamber Morphogenesis by Spatiotemporal Degradation of Cardiac Jelly. Cell Rep 23, 2455–2466.

Klaus, A., Muller, M., Schulz, H., Saga, Y., Martin, J.F., and Birchmeier, W. (2012). Wnt/beta-catenin and Bmp signals control distinct sets of transcription factors in cardiac progenitor cells. Proc Natl Acad Sci U S A 109, 10921–10926.

Kolde, R. (2019). pheatmap: Pretty Heatmaps. https://CRANR-projectorg/package=pheatmap.

Koni, P.A., Joshi, S.K., Temann, U.A., Olson, D., Burkly, L., and Flavell, R.A. (2001). Conditional vascular cell adhesion molecule 1 deletion in mice: impaired lymphocyte migration to bone marrow. J Exp Med 193, 741–754.

Langmead, B., and Salzberg, S.L. (2012). Fast gapped-read alignment with Bowtie 2. Nature methods 9, 357.

Lawrence, M., Gentleman, R., and Carey, V. (2009). rtracklayer: an R package for interfacing with genome browsers. Bioinformatics 25, 1841–1842.

Levine, M. (2011). Paused RNA polymerase II as a developmental checkpoint. Cell 145, 502–511.

Li, H., Handsaker, B., Wysoker, A., Fennell, T., Ruan, J., Homer, N., Marth, G., Abecasis, G., Durbin, R., and Genome Project Data Processing, S. (2009). The Sequence Alignment/Map format and SAMtools. Bioinformatics 25, 2078-2079.

Liao, Y., Wang, J., Jaehnig, E.J., Shi, Z., and Zhang, B. (2019). WebGestalt 2019: gene set analysis toolkit with revamped UIs and APIs. Nucleic Acids Res 47, W199–w205.

Lincoln, J., Kist, R., Scherer, G., and Yutzey, K.E. (2007). Sox9 is required for precursor cell expansion and extracellular matrix organization during mouse heart valve development. Dev Biol 305, 120–132.

Liu, H., Zhang, C.H., Ammanamanchi, N., Suresh, S., Lewarchik, C., Rao, K., Uys, G.M., Han, L., Abrial, M., Yimlamai, D., et al. (2019). Control of cytokinesis by beta-adrenergic receptors indicates an approach for regulating cardiomyocyte endowment. Sci Transl Med 11.

Liu, X., Kraus, W.L., and Bai, X. (2015). Ready, pause, go: regulation of RNA polymerase II pausing and release by cellular signaling pathways. Trends Biochem Sci 40, 516–525.

MacGrogan, D., Munch, J., and de la Pompa, J.L. (2018). Notch and interacting signalling pathways in cardiac development, disease, and regeneration. Nat Rev Cardiol 15, 685–704.

Meilhac, S.M., and Buckingham, M.E. (2018). The deployment of cell lineages that form the mammalian heart. Nat Rev Cardiol 15, 705–724.

Mikryukov, A.A., Mazine, A., Wei, B., Yang, D., Miao, Y., Gu, M., and Keller, G.M. (2021). BMP10 Signaling Promotes the Development of Endocardial Cells from Human Pluripotent Stem Cell-Derived Cardiovascular Progenitors. Cell Stem Cell 28, 96–111 e117.

Moretti, A., Caron, L., Nakano, A., Lam, J.T., Bernshausen, A., Chen, Y., Qyang, Y., Bu, L., Sasaki, M., Martin-Puig, S., et al. (2006). Multipotent embryonic isl1+ progenitor cells lead to cardiac, smooth muscle, and endothelial cell diversification. Cell 127, 1151–1165.

Munch, J., and Abdelilah-Seyfried, S. (2021). Sensing and Responding of Cardiomyocytes to Changes of Tissue Stiffness in the Diseased Heart. Front Cell Dev Biol 9, 642840.

Nagai, N., Ohguchi, H., Nakaki, R., Matsumura, Y., Kanki, Y., Sakai, J., Aburatani, H., and Minami, T. (2018). Downregulation of ERG and FLI1 expression in endothelial cells triggers endothelial-to-mesenchymal transition. PLoS Genet 14, e1007826.

Nandadasa, S., Foulcer, S., and Apte, S.S. (2014). The multiple, complex roles of versican and its proteolytic turnover by ADAMTS proteases during embryogenesis. Matrix Biol 35, 34–41.

Neuwirth, E. (2014). RColorBrewer: ColorBrewer Palettes. https://CRANR-projectorg/package=RColorBrewer.

Palpant, N.J., Pabon, L., Friedman, C.E., Roberts, M., Hadland, B., Zaunbrecher, R.J., Bernstein, I., Zheng, Y., and Murry, C.E. (2017). Generating high-purity cardiac and endothelial derivatives from patterned mesoderm using human pluripotent stem cells. Nat Protoc 12, 15–31.

Pavri, R., Zhu, B., Li, G., Trojer, P., Mandal, S., Shilatifard, A., and Reinberg, D. (2006). Histone H2B monoubiquitination functions cooperatively with FACT to regulate elongation by RNA polymerase II. Cell 125, 703–717.

Ramírez, F., Dündar, F., Diehl, S., Grüning, B.A., and Manke, T. (2014). deepTools: a flexible platform for exploring deep-sequencing data. Nucleic acids research 42, W187–W191.

Rhee, S., Paik, D.T., Yang, J.Y., Nagelberg, D., Williams, I., Tian, L., Roth, R., Chandy, M., Ban, J., Belbachir, N., et al. (2021). Endocardial/endothelial angiocrines regulate cardiomyocyte development and maturation and induce features of ventricular non-compaction. Eur Heart J 42, 4264–4276.

Robinson, M.D., McCarthy, D.J., and Smyth, G.K. (2010). edgeR: a Bioconductor package for differential expression analysis of digital gene expression data. Bioinformatics 26, 139–140.

Robson, A., Makova, S.Z., Barish, S., Zaidi, S., Mehta, S., Drozd, J., Jin, S.C., Gelb, B.D., Seidman, C.E., Chung, W.K., et al. (2019). Histone H2B monoubiquitination regulates heart development via epigenetic control of cilia motility. Proc Natl Acad Sci U S A 116, 14049–14054.

Rochais, F., Dandonneau, M., Mesbah, K., Jarry, T., Mattei, M.G., and Kelly, R.G. (2009). Hes1 is expressed in the second heart field and is required for outflow tract development. PLoS One 4, e6267.

Scharf, G.M., Kilian, K., Cordero, J., Wang, Y., Grund, A., Hofmann, M., Froese, N., Wang, X., Kispert, A., Kist, R., et al. (2019). Inactivation of Sox9 in fibroblasts reduces cardiac fibrosis and inflammation. JCI Insight 5.

Shema, E., Kim, J., Roeder, R.G., and Oren, M. (2011). RNF20 inhibits TFIIS-facilitated transcriptional elongation to suppress pro-oncogenic gene expression. Mol Cell 42, 477–488.

Shen, L., Shao, N., Liu, X., and Nestler, E. (2014). ngs.plot: Quick mining and visualization of next-generation sequencing data by integrating genomic databases. BMC Genomics 15, 284.

Shen, Y., Yue, F., McCleary, D.F., Ye, Z., Edsall, L., Kuan, S., Wagner, U., Dixon, J., Lee, L., Lobanenkov, V.V., et al. (2012). A map of the cis-regulatory sequences in the mouse genome. Nature 488, 116–120.

Skelly, D.A., Squiers, G.T., McLellan, M.A., Bolisetty, M.T., Robson, P., Rosenthal, N.A., and Pinto, A.R. (2018). Single-Cell Transcriptional Profiling Reveals Cellular Diversity and Intercommunication in the Mouse Heart. Cell Rep 22, 600–610.

Stark, R., and Brown, G. DiffBind: differential binding analysis of ChIP-Seq peak data. Bioconductor (2011).

Tian, Y., and Morrisey, E.E. (2012). Importance of myocyte-nonmyocyte interactions in cardiac development and disease. Circ Res 110, 1023–1034.

Tools, P. (2015). By Broad Institute.

VanDusen, N.J., Lee, J.Y., Gu, W., Butler, C.E., Sethi, I., Zheng, Y., King, J.S., Zhou, P., Suo, S., Guo, Y., et al. (2021). Massively parallel in vivo CRISPR screening identifies RNF20/40 as epigenetic regulators of cardiomyocyte maturation. Nat Commun 12, 4442.

Vincent, S.D., and Buckingham, M.E. (2010). How to make a heart: the origin and regulation of cardiac progenitor cells. Curr Top Dev Biol 90, 1–41.

Wang, Y., Nakayama, M., Pitulescu, M.E., Schmidt, T.S., Bochenek, M.L., Sakakibara, A., Adams, S., Davy, A., Deutsch, U., Luthi, U., et al. (2010). Ephrin-B2 controls VEGF-induced angiogenesis and lymphangiogenesis. Nature 465, 483–486.

Wickham, H. (2016). ggplot2: Elegant Graphics for Data Analysis. (New York: Springer-Verlag).

Windmueller, R., Leach, J.P., Babu, A., Zhou, S., Morley, M.P., Wakabayashi, A., Petrenko, N.B., Viatour, P., and Morrisey, E.E. (2020). Direct Comparison of Mononucleated and Binucleated Cardiomyocytes Reveals Molecular Mechanisms Underlying Distinct Proliferative Competencies. Cell Rep 30, 3105–3116 e3104.

Witzel, H.R., Jungblut, B., Choe, C.P., Crump, J.G., Braun, T., and Dobreva, G. (2012). The LIM protein Ajuba restricts the second heart field progenitor pool by regulating Isl1 activity. Dev Cell 23, 58–70.

Wu, T., Hu, E., Xu, S., Chen, M., Guo, P., Dai, Z., Feng, T., Zhou, L., Tang, W., Zhan, L., et al. (2021). clusterProfiler 4.0: A universal enrichment tool for interpreting omics data. Innovation (Camb) 2, 100141.

Xiao, Y., Hill, M.C., Zhang, M., Martin, T.J., Morikawa, Y., Wang, S., Moise, A.R., Wythe, J.D., and Martin, J.F. (2018). Hippo Signaling Plays an Essential Role in Cell State Transitions during Cardiac Fibroblast Development. Dev Cell 45, 153–169 e156.

Xiong, H., Luo, Y., Yue, Y., Zhang, J., Ai, S., Li, X., Wang, X., Zhang, Y.L., Wei, Y., Li, H.H., et al. (2019). Single-Cell Transcriptomics Reveals Chemotaxis-Mediated Intraorgan Crosstalk During Cardiogenesis. Circ Res 125, 398–410.

Yang, J., Savvatis, K., Kang, J.S., Fan, P., Zhong, H., Schwartz, K., Barry, V., Mikels-Vigdal, A., Karpinski, S., Kornyeyev, D., et al. (2016). Targeting LOXL2 for cardiac interstitial fibrosis and heart failure treatment. Nat Commun 7, 13710.

Yu, G. (2021). enrichplot: Visualization of Functional Enrichment Result. https://yulab-smutop/biomedical-knowledge-mining-book/.

Yu, G., Wang, L.G., and He, Q.Y. (2015). ChIPseeker: an R/Bioconductor package for ChIP peak annotation, comparison and visualization. Bioinformatics 31, 2382–2383.

Yuan Tang, M.H., Wenxuan Li (2016). ggfortify: Unified Interface to Visualize Statistical Result of Popular R Packages. The R Journal 8.*2*, 478–489.

Zaidi, S., Choi, M., Wakimoto, H., Ma, L., Jiang, J., Overton, J.D., Romano-Adesman, A., Bjornson, R.D., Breitbart, R.E., Brown, K.K., et al. (2013). De novo mutations in histone-modifying genes in congenital heart disease. Nature 498, 220–223.

Zhang, H., Lui, K.O., and Zhou, B. (2018). Endocardial Cell Plasticity in Cardiac Development, Diseases and Regeneration. Circ Res 122, 774–789.

Zhang, Y., Liu, T., Meyer, C.A., Eeckhoute, J., Johnson, D.S., Bernstein, B.E., Nusbaum, C., Myers, R.M., Brown, M., Li, W., et al. (2008). Model-based analysis of ChIP-Seq (MACS). Genome Biol 9, R137.

Zhao, J., Patel, J., Kaur, S., Sim, S.L., Wong, H.Y., Styke, C., Hogan, I., Kahler, S., Hamilton, H., Wadlow, R., et al. (2021). Sox9 and Rbpj differentially regulate endothelial to mesenchymal transition and wound scarring in murine endovascular progenitors. Nat Commun 12, 2564.

Zheng, G.X.Y., Terry, J.M., Belgrader, P., Ryvkin, P., Bent, Z.W., Wilson, R., Ziraldo, S.B., Wheeler, T.D., McDermott, G.P., Zhu, J., et al. (2017). Massively parallel digital transcriptional profiling of single cells. Nat Commun 8, 14049.

Zhuang, S., Zhang, Q., Zhuang, T., Evans, S.M., Liang, X., and Sun, Y. (2013). Expression of Isl1 during mouse development. Gene Expr Patterns 13, 407–412.

